# Common types of microdebris affect the physiology of reef-building corals

**DOI:** 10.1101/2023.12.04.569976

**Authors:** Jessica Reichert, Vanessa Tirpitz, Katherine Plaza, Elisabeth Wörner, Luisa Bösser, Susanne Kühn, Sebastian Primpke, Patrick Schubert, Maren Ziegler, Thomas Wilke

**Affiliations:** Department of Animal Ecology & Systematics, Justus Liebig University, Giessen, Germany; Hawaiʻi Institute of Marine Biology, University of Hawaiʻi at Mānoa, Hawaiʻi, Kāne‘ohe, USA; Department of Geoscience, University of Oslo, Oslo, Norway; Wageningen Marine Research, Den Helder, The Netherlands; Alfred-Wegener-Institute Helmholtz Centre for Polar and Marine Research, Biologische Anstalt Helgoland, Helgoland, Germany

**Keywords:** artificial clothing fibers, tire wear, secondary microplastic, marine debris, coral growth, coral photosynthesis

## Abstract

Marine debris, particularly microdebris (< 1 mm) poses a potential threat to marine life, including reef-building corals. While previous research has mainly focused on the impact of single polymer microplastics, the effects of natural microdebris, composed of a mixture of materials, have not been explored. Therefore, this study aimed to assess the effects of different microdebris, originating from major sources of pollution, on reef-building corals. For this, we exposed two scleractinian coral species, *Pocillopora verrucosa* and *Stylophora pistillata*, known to frequently ingest microplastics, to four types of microdebris in an 8-week laboratory experiment: fragmented environmental plastic debris, artificial fibers from clothing, residues from the automobile sector consisting of tire wear, brake abrasion, and varnish flakes, a single polymer microplastic treatment consisting of polyethylene particles, and a microdebris-free control treatment. Specifically, we (I) compared the effects of the different microdebris on coral growth, necrosis, and photosynthesis, (II) investigated the difference between the microdebris mixtures and the exposure to the single polymer treatment, and (III) identified potential mechanisms causing species-specific effects by contrasting the feeding responses of the two coral species on microdebris and natural food. We show that the fibers and tire wear had the strongest effects on coral physiology, with *P. verrucosa* being more affected than *S. pistillata*. Both species showed increased volume growth in response to the microdebris treatments, accompanied by decreased calcification in *P. verrucosa*. Photosynthetic efficiency of the symbionts was enhanced in both species. The species-specific physiological responses might be attributed to feeding reactions, with *P. verrucosa* responding significantly more often to microdebris than *S. pistillata*. These findings highlight the effect of different microdebris on coral physiology and the need for future studies to use particle mixtures to better mimic naturally occurring microdebris and assess its effect on corals in more detail.

**Highlights:**

- The effects of major sources of microdebris pollution on coral physiology were compared.
- Overall, microdebris had only minor impacts on coral physiology.
- Artificial fibers and tire wear caused the strongest effect on coral physiology.
- Single polymer and complex microdebris caused similar, yet species-specific effects.
- Species-specific effects might be due to different feeding behaviors.

**Graphical abstract:** 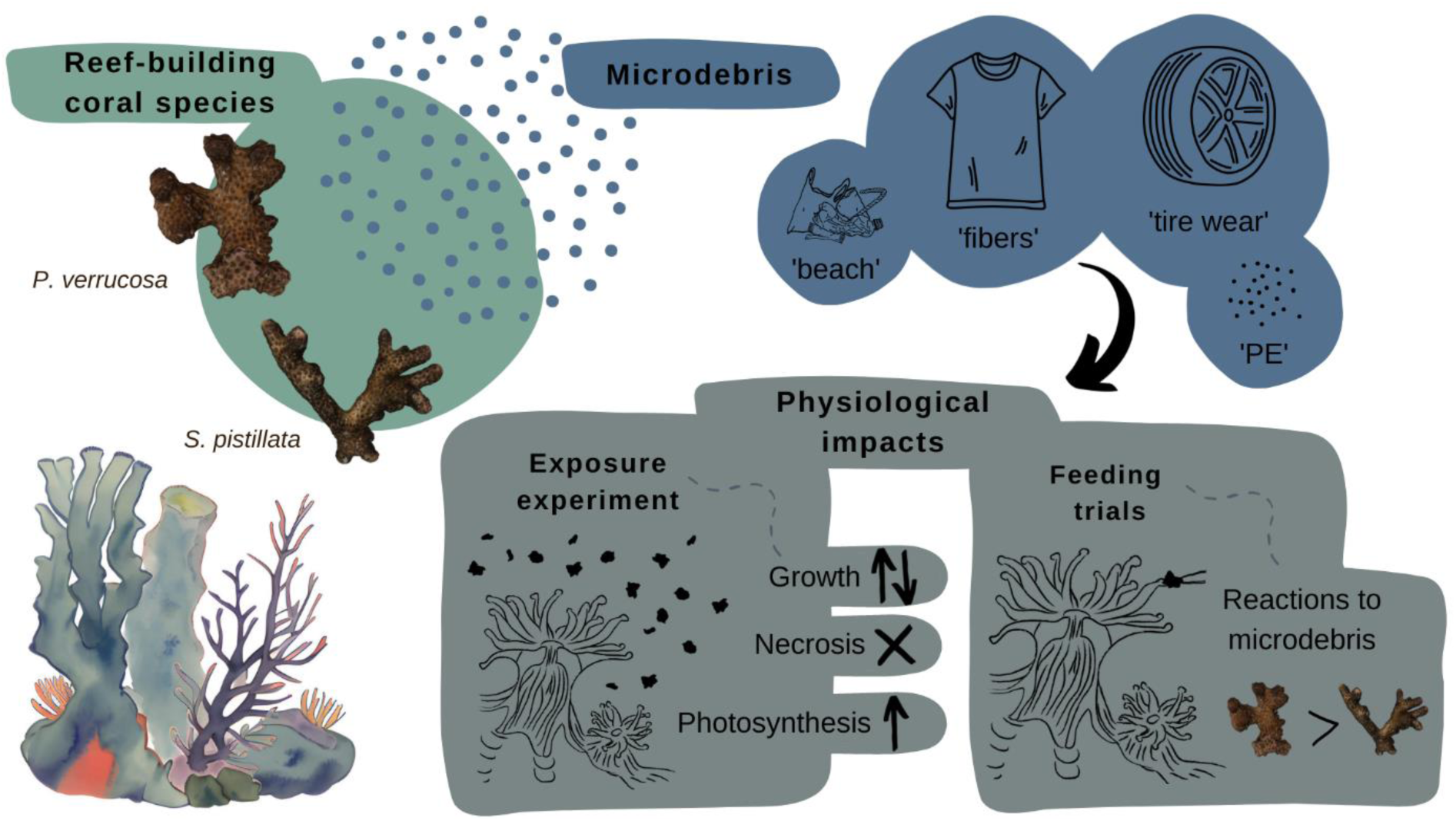

## 1 Introduction

Since the expansion of organic chemistry in the 1950s, the pollution of the world’s oceans with marine debris – manufactured or processed material that is discarded in marine and coastal environments (UNEP, 2009) – has become a tremendous problem (Gregory, 2009; Provencher et al., 2017; Ryan, 2018; Wilcox et al., 2018). Small particles (i.e., microdebris) in particular pose a threat to marine life, as they are frequently ingested by mollusks, fish, or corals (Ding et al., 2022; Huang et al., 2021; Savoca et al., 2021). Tire wear, artificial fibers from clothing, and discarded plastic products are major contributors to pollution (Carr, 2017; Coyle et al., 2020; Sommer et al., 2018; Wagner et al., 2018). Yet, little is known about the effect of microdebris on reef-building corals, the main framework builders of coral reef ecosystems.

Most studies to date have shown the adverse influence of single polymer types, often in the form of unfouled manufactured beads (Barnes et al., 2009; Paul-Pont et al., 2018; Phuong et al., 2016). Yet, marine microdebris (i.e., particles between 1 µm and 1 mm in size) occurs as a mixture in a variety of types and shapes (Galgani et al., 2015; Hartmann et al., 2019; Kroon et al., 2018; UNEP, 2009). The most obvious source of microdebris is plastic litter of everyday products that enters the ocean and degrades into smaller particles over time, consistent with our classic understanding of secondary microplastics (Andrady, 2011; Hidalgo-Ruz et al., 2012). It includes polyethylene (PE), polypropylene (PP), polyethylene terephthalate (PET), polyamide (PA), and polyvinyl chloride (PVC), with PE being one of the most common particle types found (Andrady, 2011; Kühn et al., 2018). Another major source of marine microdebris is artificial fibers from clothing, which are released during the washing process and mainly consist of polyester, acrylic, PP, PE, and PA (Boucher and Friot, 2017; Browne et al., 2011; Siegfried et al., 2017). Traffic-related residues, such as tire wear or break abrasion, also contribute substantially to marine microdebris, but are often underestimated or omitted in monitoring approaches, due to their inconsistent classification as microplastic (Jeong et al., 2022; Rødland et al., 2022; Siegfried et al., 2017).

Reef-building corals – the ecosystem engineers of coral reefs – have been found to be affected by microplastics (reviewed in Huang et al., 2021). Exposure to microplastics is often associated with subtle but negative effects on growth and calcification (Chapron et al., 2018; Hankins et al., 2021; Reichert et al., 2019). Occasionally, compromised coral health, such as bleaching or necrosis, is observed after prolonged exposure or under high pollution scenarios (Reichert et al., 2019, 2018; Syakti et al., 2019). Further, microplastic exposure has been found to impact the associated photosymbionts, affecting their photosynthetic rate and efficiency (Chapron et al., 2018; Hankins et al., 2021; Mendrik et al., 2021; Reichert et al., 2019). The physiological impacts are often attributed to the handling and ingestion of particles, which has been frequently observed (Allen et al., 2017; Hall et al., 2015) and is suspected to affect their feeding behavior (Chapron et al., 2018; Savinelli et al., 2020). These effects on feeding coincide with elevated energetic needs, caused by mucus production for cleaning from the particles, as well as overall increased stress and immune responses of the coral (Chen et al., 2022; Lanctôt et al., 2020; Reichert et al., 2019; Tang et al., 2018). However, the effects of microplastics are species-specific and some species, such as *Porites* spp. or *Stylophora* spp. appear to be less affected than others, such as *Pocillopora* spp. or *Acropora* spp. (Mendrik et al., 2021; Mouchi et al., 2019; Reichert et al., 2019). Reasons for the species-specific differences are still largely unknown but might be attributed to differential feeding behavior, particle recognition, or colony shape (Hankins et al., 2022; Martin et al., 2019; Rades et al., 2022; Reichert et al., 2022).

Observations on the effects of microplastics on reef-building corals have mostly been made under controlled exposure to particles of a single plastic polymer, with standardized shape and size. Yet, little is known about how the effects of microplastics of individual polymers on reef-building corals compare to the effects of microdebris, which is found as a complex mixture in natural environments (reviewed in Huang et al., 2021). Thus, the major goal of this study was to assess the effect of microdebris, originating from major sources of ocean pollution, on reef-building corals. These mixtures comprised fragmented plastic debris from the beach, artificial fibers from clothing, and residues from the automobile sector, consisting of tire wear, brake abrasion, and varnish flakes. We compared the effects of microdebris exposure to a single-type microplastic treatment consisting of polyethylene and a microdebris-free control treatment. Two scleractinian coral species (*Pocillopora verrucosa* (Ellis & Solander, 1786) and *Stylophora pistillata* (Esper, 1792)) were used as test organisms. These ecologically similar species both belong to the Pocilloporidae family, but are known to be differentially affected by microplastics in their physiology (Lanctôt et al., 2020; Reichert et al., 2021, 2018). Since impacts on coral physiology might be linked to particle detection and handling, influencing the frequency of microplastic ingestion and, consequently, the overall effects, we investigate the feeding responses of the corals to the tested microdebris.

Specifically, we (I) compared the effects of different microdebris mixtures on coral growth, necrosis, and photosynthesis, (II) investigated the difference between the microdebris mixtures and the exposure to the single polymer treatment, and (III) identified potential mechanisms causing species-specific effects by contrasting the feeding responses of the two reef-building coral species on microdebris and natural food.

## 2 Material and Methods

### 2.1 Experimental design and replication

The study assessed the effects of different types of common microdebris on the two scleractinian coral species *Pocillopora verrucosa* and *Stylophora pistillata* (experimental overview: Figure 1). These species both have small corallite sizes (*P. verrucosa*: 0.3-0.7 mm and *S. pistillata*: 0.9-1.4 mm), are presumed to be sensitive to environmental stressors, and have been shown to frequently interact with and regularly ingest microplastics (Lanctôt et al., 2020; Reichert et al., 2018). The microdebris treatments were selected to represent major sources of pollution (Agamuthu et al., 2019). These are secondary marine microplastics derived from fragmented plastic debris from the beach (treatment: ‘beach’), an assortment of artificial fibers from clothing (treatment: ‘fibers’), residues from the automobile sector consisting of tire wear, brake abrasion, and varnish flakes (treatment: ‘tire wear’), and a single polymer exposure with polyethylene (treatment: ‘PE’). The average size of the microdebris varied depending on the treatment within a range of 350 to 1000 µm, which is within the size range for particles commonly consumed by corals (Houlbrèque and Ferrier-Pagès, 2009).

**Figure 1:**
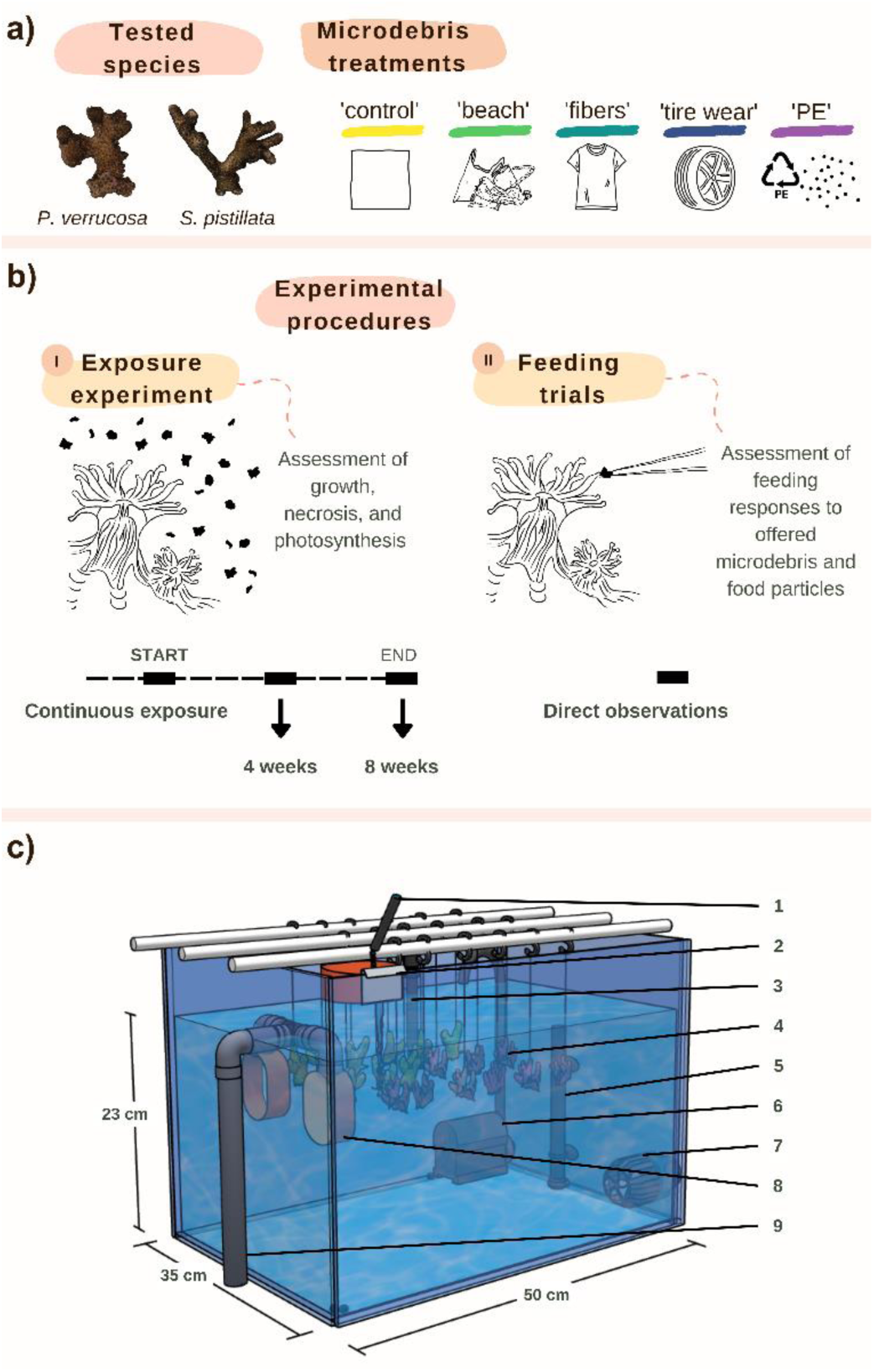
Experimental design. a) The study tested the effects of the microdebris treatments ‘beach’, ‘fibers’, ‘tire wear’, and the single polymer treatment ‘PE’ on the two reef-building coral species *Pocillopora verrucosa* and *Stylophora pistillata*. b) The effects on growth, necrosis, and photosynthesis were assessed after 4 and 8 weeks of an exposure experiment (I). The feeding responses to the different particle types were studied in feeding trials (II). c) The exposure experiment was conducted in 15 recirculating 40 L tanks. Six fragments per species originating from different colonies were placed in each tank. The 3D model illustrates the technical setup that allowed for continuous water circulation while retaining particles within the tanks: 1. Water inflow; 2. Inflow filter; 3. Temperature sensor; 4. Hanging coral fragments; 5. Heater; 6. Turnover pump; 7. Wavemaker pump; 8. Outflow filters; 9. Water outflow.

Physiological responses of corals to the microdebris treatments were examined in an 8-week exposure study. The experiment was conducted in 15 tanks, each with a volume of 40 L (n = 3 tanks per treatment). For this, six mother colonies were used per species (colony origin and CITES numbers are given in Table S1). One fragment from each colony was placed in each of the 15 tanks resulting in n = 18 replicates per treatment and species.

Feeding responses of the tested coral species to the four types of microdebris (beach, fibers, tire wear, and PE) as well as copepods, serving as natural control feed, were assessed in direct feeding trials, performed in separate feeding chambers under a stereomicroscope. Fifty feeding responses were assessed in one fragment from each of the six colonies, resulting in n = 300 feeding offers per particle type and species.

### 2.2 Coral fragment preparation

Corals were fragmented 6–9 weeks before the experiment to allow for tissue recovery in the cutting areas. Corals were cut from terminal branches into fragments of 3–5 cm and tied to fishing lines to avoid the buildup of detritus on the colonies and decrease algal overgrowth. To control for position effects, replicates from the same colony were placed in the same spot in each tank.

### 2.3 Technical setup and maintenance

This experiment was conducted in the Ocean2100 facility at Justus Liebig University Giessen. The experimental tanks (n = 15, Figure 1) were part of a closed laboratory setup, containing 2,300 L artificial seawater (salt: Instant Ocean, Aquarium Systems, France). The setup contained a technical tank, fitted with a protein skimmer and a calcium reactor and other tanks harboring reef organisms. The experimental tanks were filled with 38 L of seawater (salinity: 35) and connected to the system at an exchange rate of 78 L h^-1^. The outflows of the tanks were equipped with two self-made 65-µm filters to retain the microdebris within the tanks and allow for a biofilm to form on the particles. Incoming water was filtered before entering the tanks through a mesh (mesh size: 65 µm) to minimize contamination through algae or detritus as well as potential cross-contamination with microdebris from the connected system.

Light was provided at an intensity of 120 µmol photons m^-2^ s^-1^ (Apogee Quantum Flux MP-200, Apogee Instruments, USA; measuring range: 410–655 nm, PAR) with a 10:14 h light:dark photoperiod (four SunaECO LED Lighting Strips per tank, Tropical Marine Centre, United Kingdom). Water flow in the tanks was created with a wavemaker pump generating horizontal pulsating currents (easySTREAM pro Wavemaker Pump, ES-18, Aqualight GmbH, Germany) and a turnover pump (Submarine Water Pump, Resun S-700, Resun, China) generating vertical currents to submerge floating debris, which resulted in a flow velocity of 3.8 cm s^-1^ (OTT MF pro, HydroMet, Germany). Water temperatures were maintained at 27 ± 0.3 °C using feedback-controlled titanium heaters (Schego Heater 300 W, Protection class IP 68, SCHEGO Schemel & Goetz GmbH & Co KG, Germany) coupled to temperature sensors (GHL Temp Sensor digital, GHL Advanced Technology GmbH & Co. KG, Germany) and GHL aquarium computers (ProfiLux 3 and ProfiLux 4, GHL Advanced Technology GmbH & Co. KG, Germany).

Gastropods (12 x *Turbo* sp. and 2 x *Euplica* sp.) were included in each tank to reduce algal growth. Corals were fed daily with copepods (∼0.5 g per tank, Calanoide Copepoden, Zooschatz, Germany) and amino acids were added to the water column (∼0.5 mL amino acid mixture per tank, Pohl’s Xtra special, Korallenzucht.de Vertriebs GmbH, Germany). All maintenance activities were carried out so that microdebris remained in the tank, e.g., if filters were cleaned, they were rinsed above the tanks before being removed. Further details on the technical setup and maintenance can be found in Supplementary Material and Methods.

### 2.4 Microdebris exposure treatments

To represent secondary marine microplastics, plastic debris was used (treatment: ‘beach’; Figure 2a). The plastic was collected on a beach in Texel, The Netherlands, and fragmented by cryogenic milling, resulting in a representative mix of 60.9 % polyethylene, 27.7 % polypropylene, 3.1 % polyamide, 1.9 % polyurethane, 1.7 % polystyrene, 0.9 % polyvinyl chloride, 0.5 % polyethylene terephthalate, 0.2 % ethylene-vinyl acetate, 0.1 % ethylene-vinyl acetate, and 3.1 % unidentified particles (% by mass, details on the collected mixture are provided in Kühn et al., 2018). The particles were sieved to a particle size > 100 µm. The upper limit was maintained and not further sieved, due to the limited amount of debris available, resulting in a mean particle length of 346 ± 240 µm and a mean particle width of 184 ± 126 µm.

**Figure 2:**
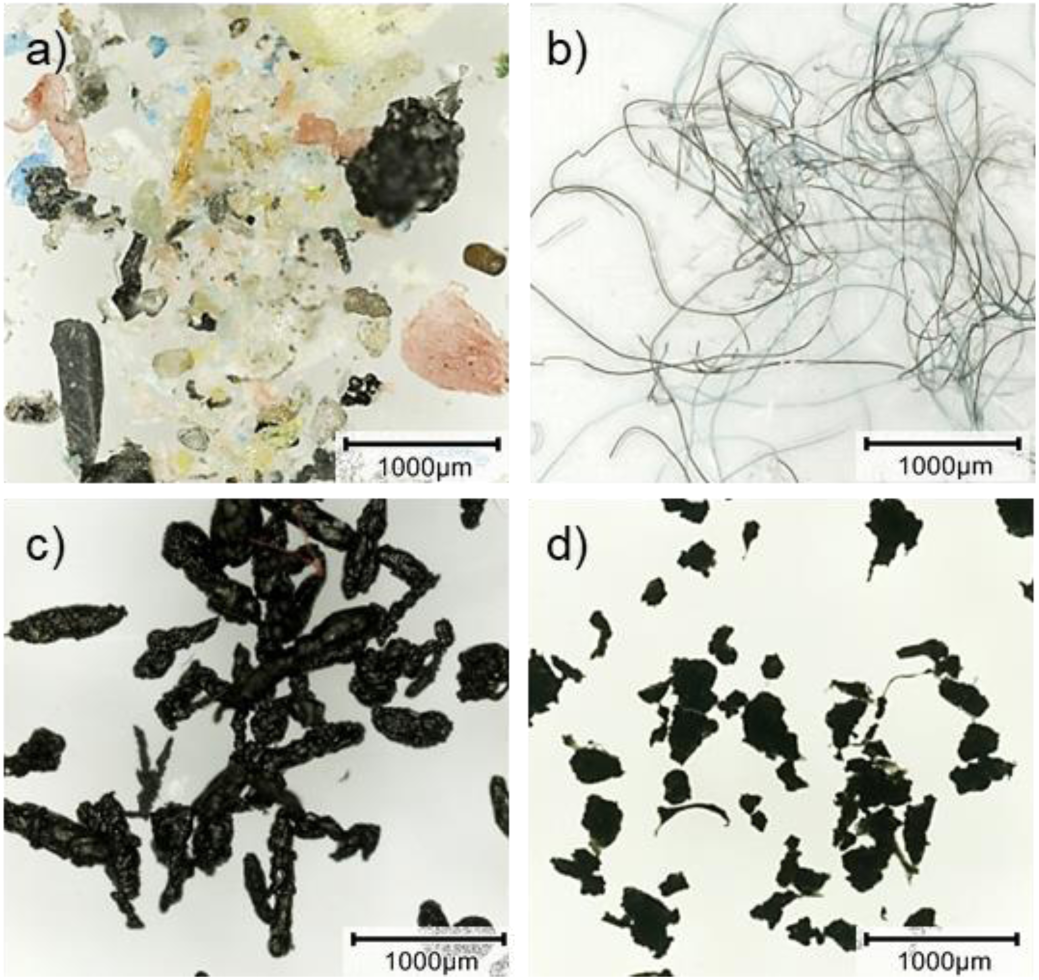
Types of microdebris used in the experiment: a) Secondary marine microplastics, composed of fragmented plastic debris (treatment: ‘beach’), b) artificial fibers from clothing (treatment: ‘fibers’), c) residues from the automobile sector, consisting of tire wear, brake abrasion, and varnish flakes (treatment: ‘tire wear’), and d) polyethylene particles (treatment: ‘PE’).

To represent a naturally occurring mixture of fibers, synthetic fabrics were used (treatment: ‘fibers’; Figure 2b). Fibers were produced from clothing, i.e., a grey pullover consisting of 70 % acrylic and 30 % polyamide as well as a black and a green fleece, both made of 100 % polyester, manually cut into small pieces, resulting in a mean length of 1,006 ± 840 µm and a mean width of 16 ± 4 µm. The fibers were mixed to consist of 67 % Polyester, 23 % acrylic, and 10 % polyamide to mimic natural occurrences in the ocean (Browne et al., 2011).

To represent residues from the automobile sector, tire wear particles, brake abrasion, and varnish flakes were swept from a cart track (treatment: ‘tire wear’; Figure 2c). Carts were at most 12 months old, built with a hydraulic brake system, and equipped with wheels from Dunlop (Cart Tires, Goodyear Dunlop Tires Germany GmbH, Germany). Particles were strained with a metal sieve (stainless steel ISO 3310-1, Joachim Edinger Industrievertretungen, Germany) to 100–355 µm, resulting in a mean particle length of 583 ± 235 µm and a mean particle width of 218 ± 71 µm.

To represent a treatment commonly used in previous experimental approaches, a single polymer reference with polyethylene (PE) microplastic particles was included (treatment: ‘PE’; Figure 2d), which is one of the most common polymers in the marine environment (Nie et al., 2019; Patterson et al., 2020; Saliu et al., 2018). Black polyethylene particles of irregular shape (Novoplastik Produktions-und Vertriebs-GmbH, Germany) were used as in previous studies (Reichert et al., 2019, 2018). Particles were strained to 100–355 µm, resulting in a mean particle length of 371 ± 112 µm and a mean particle width of 226 ± 70 µm. Although microdebris sizes differ significantly depending on the treatment, their mean sizes, ranging between 350 and 1000 µm, all falls into the size spectrum corals commonly feed on (Houlbrèque and Ferrier-Pagès, 2009).

### 2.5 Microdebris preparation and concentration

Particles were incubated for at least two days in seawater to initiate biofilm formation on the surface and ease submerging in the water. For this, 0.4 g of particles were incubated with 10 mL of seawater in 15 mL falcon tubes for each tank. The exposure was standardized by mass and the incubated particles were added to each tank, equivalent to ∼10 mg per L and resulting in a target concentration of ∼200 particles per L in each treatment. Microdebris concentrations were monitored and maintained over the course of the experiment, by adding particles when necessary (e.g., due to debris removal during tank maintenance, ingestion, and overgrowth by the corals). The concentration of microdebris in each tank was determined once a week. For this, 100 mL of water was taken with a dropper pipette from the water surrounding the corals (n = 3 per tank). Each sample was vacuum filtered onto a separate 8 µm cellulose filter (Whatman filter papers Grade 540, General Electric Healthcare Life Sciences, United States). Debris was counted under a stereomicroscope and extrapolated to assess the concentration as particles per L. Although water flow in the tanks was optimized to distribute all debris types equally in the water column, they exhibited different buoyancy and behavior in the water (e.g., tire wear and beach particles tended to sink to the bottom, while fibers and PE particles rather accumulated at the surface). This resulted in slightly different debris concentrations (particles per L) in the water column, depending on the treatment (beach: 86 ± 63, fibers: 228 ± 206, tire wear: 66 ± 50, and PE: 242 ± 126 particles per L, mean ± SD, Figure S1). Fibers and PE polymer identities were confirmed by FTIR (Figure S2). The particle composition of the beach treatment is characterized by Kühn et al. (2018). Identification of tire wear particles could not be included due to technical limitations.

### 2.6 Growth measurements and necrosis

To study the physiological response of the coral host to exposure to the different microdebris, growth in tissue surface area, skeletal volume, and calcification was measured. Tissue surface area and volume of the coral fragments were assessed at the beginning, and after 4 and 8 weeks of the experiment via 3D scanning (Artec Spider 3D with Artec Studio 11, Artec Europe, Luxembourg, settings see Table S2) following established procedures (Reichert et al., 2016). Briefly, fragments were mounted on toothpicks and scanned in air in two full rotations with the handheld scanner. The sensitivity of the scanner was set at 60 %. For tissue surface area determination, bleached or necrotic tissue was removed from the model, so that only healthy coral surface remained. To study coral tissue necrosis, the difference between the total surface area of each coral fragment and its living tissue surface area was calculated and expressed as percentage of the total tissue surface area. Models were exported as .obj files and surface area and volume were calculated in MeshLab Visual Computing Lab-ISTI-CNR (v.1.3.4 Β, 2014) using the ‘compute geometric measures’ tool. Coral calcification was assessed using buoyant weight measurements (Davies, 1989). For this, coral fragments were weighed with a precision balance (Kern KB 360-3N, KERN & Sohn GmbH, Germany, precision: 0.001 g) in a separate 18 L tank. The change in buoyant weight was converted to dry weight using an aragonite density of 2.93 g cm^-3^.

### 2.7 Photosynthesis and respiration of the coral holobiont

Rates of photosynthesis and respiration of the coral holobiont were measured in individual temperature-controlled incubations, conducted in glass jars with 1,050 mL water volume (Weck, Germany). Coral fragments were processed colony-wise (n = 15 fragments from each species) together with six control incubations without coral fragments. Oxygen was measured before and after the incubations using an optical oxygen probe (FDO 925-3) coupled to a multiparameter probe (Multi 3620 IDS SET G, WTW, Germany). Net photosynthesis was measured in light (162 ± 9 µmol photons m^-2^ s^-1^) during daytime (2 hours into the photoperiod). Respiration was measured after light incubations in darkness. Incubations (both light and dark) lasted 90 min after 4 weeks and 60 min after 8 weeks, to account for the increased fragment size and avoid oxygen saturation. Differences in oxygen values were normalized to water volume and the duration of the incubation, and corrected for photosynthesis/respiration in the water by subtracting mean measurements of the control incubations. Gross photosynthesis was calculated as the sum of net photosynthesis and respiration values.

### 2.8 Photosynthetic efficiency of symbionts

The photosynthetic efficiency of the photosymbionts was determined using a Pulse-Amplitude-Modulation fluorometer (PAM, Ralph and Gademann, 2005) at the beginning and after 4 and 8 weeks of the experiment. Data was collected with the ‘PAM-2500 Portable Chlorophyll Fluorometer’ (Walz GmbH, Germany) 5 mm above and at an angle of 60° to the coral tissue using a distance clip. PAM settings were set to measure stable fluorescence levels (F, Table S3). Effective photosynthetic efficiency (ΔF/Fm’) was measured for each fragment during daytime (6 hours into the photoperiod, n = 3 per fragment). Maximum photochemical efficiency (Fv/Fm) was measured after dark acclimation at the end of the day (after 40 minutes of darkness, n = 1 per fragment). Rapid light curves (RLC) were generated to assess the maximum relative electron transport rate (rETR_max_), the minimum saturating irradiance (E_k_), and the efficiency of light capture in the light-limited phase (α). The RLCs consisted of successive measurements of the effective quantum yield (Fv/Fm) under ambient light, where light intensity increased in 11 steps (0, 4, 29, 99, 196, 361, 617, 979, 1,384, 1,661, 2,013 µmol photons m^-2^ s^-1^) lasting 10 sec each. Hyperbolic tangent functions (Jassby and Platt, 1976), which included an exponent for photoinhibition at high irradiances (Platt et al., 1980), were fitted to each curve to extract rETR_max_, E_k_, and α. RLCs were measured during daytime (3 hours after light until 2 hours before darkness).

### 2.9 Feeding response to microdebris

Feeding responses of the coral fragments to the four types of microdebris (beach, fibers, tire wear, and PE) as well as copepods as a natural control feed were assessed in additional coral fragments that were not included in the tank experiment (n = 6 per species). Coral fragments were cut from the six colonies as described above and mounted to self-made concrete bases to ease the handling in the feeding trials. Particles were manually offered to the coral fragments in individual feeding chambers (200 mL) under a stereomicroscope (Leica Microsystems GmbH, Leica EZ4, Germany). The microdebris was sampled from the treatment tanks to provide particles covered with a biofilm. The natural control feed (Calanoide Copepoden, Zooschatz, Germany) was defrosted in seawater for 1 h before feeding. Five coral polyps were consecutively fed per fragment in 10 repetitions conducted over two days, resulting in n = 300 feeding trials per treatment per species. Using forceps, the particle was placed in contact with a tentacle for five seconds and the responses to, the uptake, and egestion of the particle were assessed. Reactions were classified as positive if the particle remained attached to the coral tentacle and the time of the reaction was recorded. Reactions were classified as negative if the particle was not retained and released. Particle uptake was classified as positive if the particle was completely ingested by the coral mouth. Coral polyps that ingested particles were observed for up to 2 h to record a potential egestion, which was defined as complete upon release from the tentacles and mesenterial filaments. The time the particle was ingested was noted accordingly.

### 2.10 Statistical analyses

We performed all data processing and analyses in the R statistical environment (v.3.6.1; R Core Team, 2019). Differences in physiological responses between the microdebris treatments (beach, fibers, tire wear, and PE) and the control were assessed for each species (*P. verrucosa* and *S. pistillata*) and timepoint (after 4 and 8 weeks) using linear mixed-effects models (for surface area growth, volume growth, calcification, net photosynthesis, respiration, gross photosynthesis, effective quantum yield (Y(II)), maximum quantum yield (Fv/Fm), maximum relative electron transport rate (rETR_max_), minimum saturating irradiance (E_k_), and efficiency of light capture (α)) and generalized mixed effects models (for necrotic tissue surface area). Models were constructed with treatment as a fixed factor and colony ID set as a random factor, on scale- or log-transformed data, followed by holm-adjustment for multiple testing. Details on the model specification are given in Tables S4–6. Differences in physiological responses between the microdebris mixtures (beach, fibers, tire wear) and the single polymer PE were assessed for each species over both time points using linear mixed-effects models. Models were constructed with treatment as fixed factor and timepoint and colony ID set as a random factor, on scale or log-transformed data, followed by holm-adjustment for multiple testing. Differences in polyp behavior (i.e., reaction, uptake, and egestion) between the microdebris treatments (beach, fibers, tire wear, and PE) and natural food (copepods) for each species were tested with pairwise Fisher’s exact tests, followed by holm-adjustment for multiple testing. Differences in interaction times between microdebris types and food in cases of positive reactions, uptake events, and rejection events were tested with Wilcox rank-sum tests. Differences in coral feeding behavior (i.e., reactions, uptake, and egestion) between *Pocillopora verrucosa* and *Stylophora pistillata* to the different microdebris treatments (beach, fibers, tire wear, and PE) and natural food (copepods) were tested with Fisher’s exact tests.

## 3 Results

### 3.1 Effects of microdebris mixtures on growth rates and necrosis

The microdebris mixtures differed in their effects on coral growth rates (Figure 3a, b, Figure S3, Table S4, linear mixed-effects models). While most microdebris treatments had no significant impacts after 4 and 8 weeks, both enhancing and decreasing effects on growth rates were observed occasionally. Volume growth of *P. verrucosa* was three-fold higher in the fibers treatment after 8 weeks of exposure than in the control (fibers: 0.012 ± 0.011 cm^3^ cm^-2^ vs. control: 0.004 ± 0.012 cm^3^ cm^-2^, p = 0.006). In *S. pistillata* volume growth in the PE treatment was 36 % higher after 4 weeks of exposure than in the control (PE: 0.03 ± 0.029 cm^3^ cm^-2^ vs. control: 0.022 ± 0.028 cm^3^ cm^-2^, p = 0.050). Calcification of *P. verrucosa* in the beach and the fibers treatment decreased by 40 % and 32 %, respectively, compared to the control, after 8 weeks of exposure (beach: 8.59 ± 7.80 mg cm^-2^ and fibers: 9.71 ± 7.62 vs. control: 14.2 ± 9.55 mg cm^-2^, p = 0.004 and p = 0.026, respectively, Figure 3b). Calcification of *S. pistillata* was not affected and the microdebris treatments had no effects on growth in surface area at any time point (Figure S3, p > 0.05). Necrosis was not affected significantly (Figure 3c).

**Figure 3:**
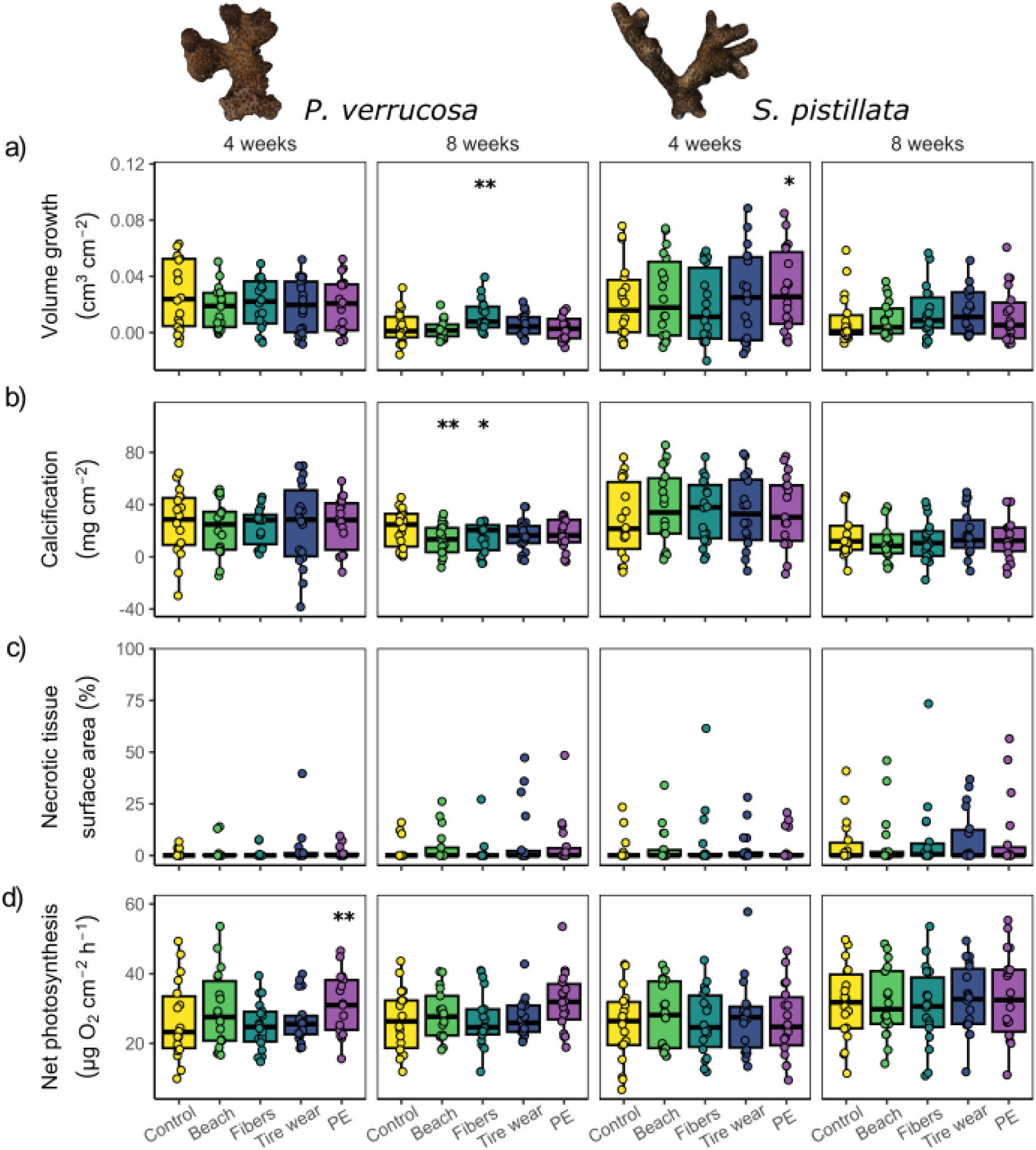
Effects of microdebris treatments (beach, fibers, tire wear, and PE) compared to the control (no microplastics added) on coral growth (volume, and calcification), necrosis, and net photosynthesis for *P. verrucosa* and *S. pistillata* after 4 and 8 weeks. Data are displayed as box-and-whisker plots with raw data points; lines indicate medians, boxes indicate the first and third quartile, and whiskers indicate ± 1.5 IQR. Significant differences between the control and the microdebris treatments are derived from linear mixed effects models or generalized linear mixed effects models followed by holm-adjusted posthoc comparison (n = 18) and are defined as p < 0.001 (***), p < 0.01 (**), and p < 0.05 (*).

### 3.2 Changes in photosynthesis and respiration

Most microdebris treatments had no significant effect on the photosynthetic productivity of the tested coral species (Figure 3d, Figure S4, Table S5, linear mixed-effects models). Only net and gross photosynthesis of *P. verrucosa* were 22 % and 15 % higher in the PE treatment after 4 weeks in comparison to the control (PE: 32.7 ± 11.7 µg O_2_ cm^-2^ h^-1^ vs. control: 26.6 ± 11.2 µg O_2_ cm^-2^ h^-1^, p = 0.005 and PE: 46.8 ± 13.4 µg O_2_ cm^-2^ h^-1^ vs. control: 39.9 ± 11.7 µg O_2_ cm^2^ h^-1^, p = 0.015, respectively). The effects were similar, but less pronounced after 8 weeks (p = 0.052 and p = 0.064, respectively). Respiration was also not affected significantly (Figure S4, p > 0.05).

### 3.3 Changes in photosynthetic efficiency

More impacts were observed on parameters relating to the photosynthetic efficiency of the associated photosymbionts of *P. verrucosa* and *S. pistillata*, which was enhanced by the exposure to microdebris (Figures 4, S5, S6, and S7, Table S6, linear mixed-effects models). For *P. verrucosa*, microdebris caused small but significant increases of ∼2 % in effective quantum yield (Y(II); beach: 0.615 ± 0.034, p = 0.025, fibers: 0.614 ± 0.029, p = 0.025, and tire wear: 0.618 ± 0.029, p = 0.002 vs. control: 0.604 ± 0.029) after 8 weeks of exposure. Similarly, the efficiency of light capture was enhanced (α; beach: 0.244 ± 0.019, p = 0.048, fibers: 0.245 ± 0.025, p = 0.035, and tire wear: 0.245 ± 0.018, p = 0.035 vs. control: 0.234 ± 0.022) after 8 weeks of exposure. Further, the fibers treatment caused an increased relative electron transport rate (rETR_max_; fibers: 122 ± 25.2 vs. control: 101 ± 21.2, p = 0.006) and reduced minimum saturating irradiance (E_k_; fibers: 499 ± 93.3 vs. control: 429 ± 78.9, p = 0.015) after 8 weeks of exposure. In contrast, fewer parameters of the photosynthetic efficiency of the photosymbionts of *S. pistillata* were affected by the exposure to microdebris. The exposure to tire wear caused an increase of ∼4 % in effective quantum yield both after 4 and 8 weeks of exposure (Y(II); tire wear: 0.607 ± 0.033, p = 0.035 vs. control: 0.585 ± 0.035, p < 0.001 and tire wear: 0.602 ± 0.037 vs. control: 0.58 ± 0.035, p = 0.010, respectively). The exposure to fibers caused an increase of ∼3 % in effective quantum yield after 4 weeks of exposure (fibers: 0.600 ± 0.032 vs. control: 0.585 ± 0.035, p = 0.030), which was no longer evident after 8 weeks (p = 0.766).

**Figure 4:**
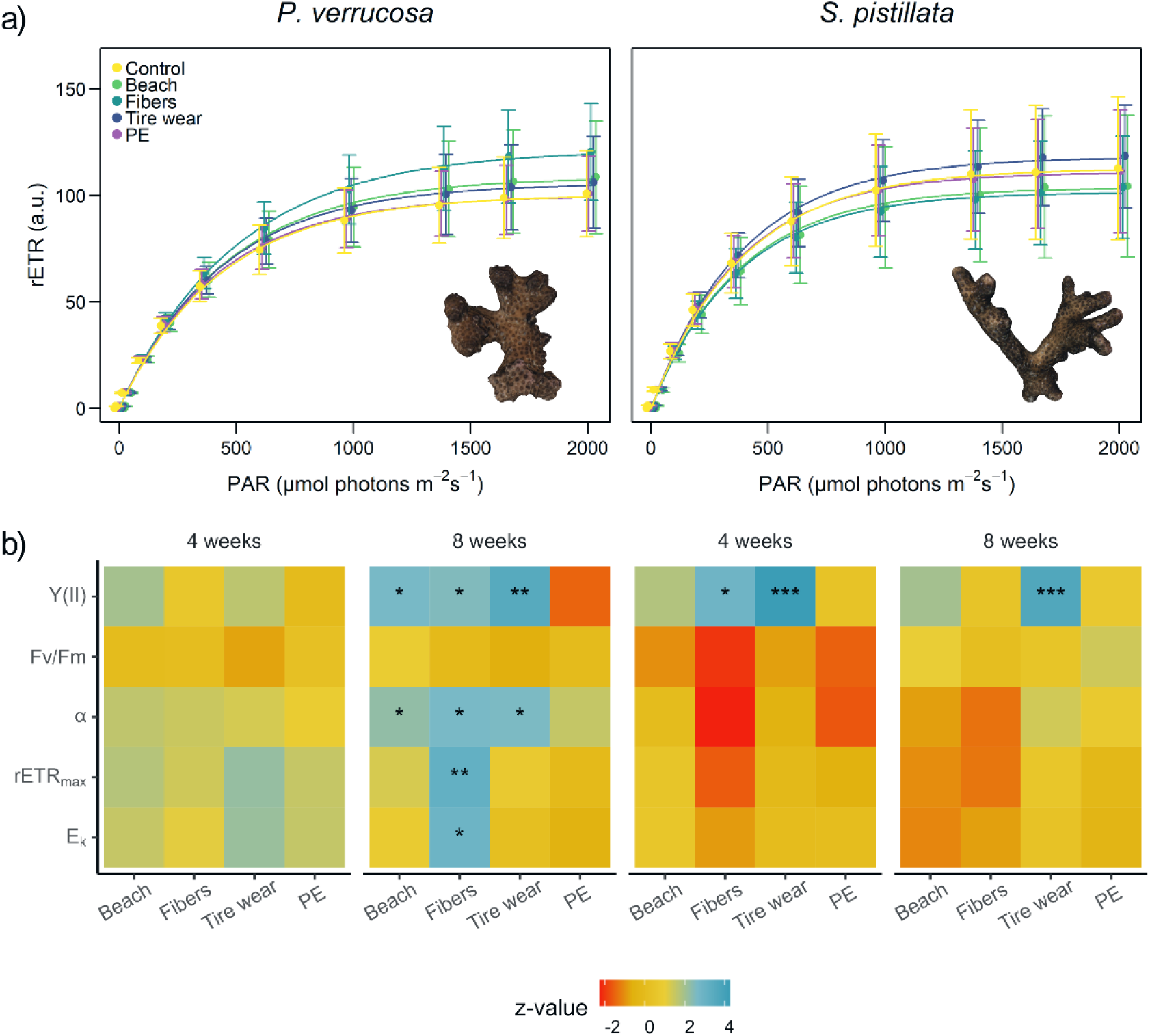
Photosynthetic efficiency of *P. verrucosa* and *S. pistillata* after 8 weeks of exposure to the different microdebris treatments (beach, fibers, tire wear, PE) and a microplastic-free control. a) Rapid light curves constructed from rETR measured at increasing PAR intensities after 8 weeks of exposure. b) Effects of microdebris treatments in comparison to the control on photosynthetic efficiency (effective quantum yield: Y(II), maximum quantum yield: Fv/Fm, maximum relative electron transport rate: rETR_max_, minimum saturation irradiance: E_k_, and efficiency of light capture: α) of *P. verrucosa* and *S. pistillata* after 4 and 8 weeks. Statistical differences (z-value) between the control and the microdebris treatments are illustrated as heatmap with higher values (blue) indicating positive effects and lower values (red) indicating negative effects compared to the control. Significant differences are marked with asterisks, defined as p < 0.001 (***), p < 0.01 (**), and p < 0.05 (*), and derived from linear mixed-effects models followed by holm-adjusted posthoc comparison (n = 18).

### 3.4 Difference between single polymer and complex microdebris treatments

Overall, the effects of the single polymer treatment PE on the tested 12 physiological response variables of the corals did not differ significantly from those of the three complex microdebris treatments (Figure 5a, Permutational Multivariate Analysis of Variance, p = 0.615, Table S7). Comparing the effects of the PE treatment to the three complex microdebris treatments on the individual physiological parameters reveals parameter- and species-specific differences. PE had less pronounced effects on effective quantum yield than the other microdebris treatments, but stronger effects on net photosynthesis (Figure 5b, Table S8, Linear mixed effects models). Further, the differences occurred mainly in *P. verrucosa*, and were less pronounced for *S. pistillata*.

**Figure 5:**
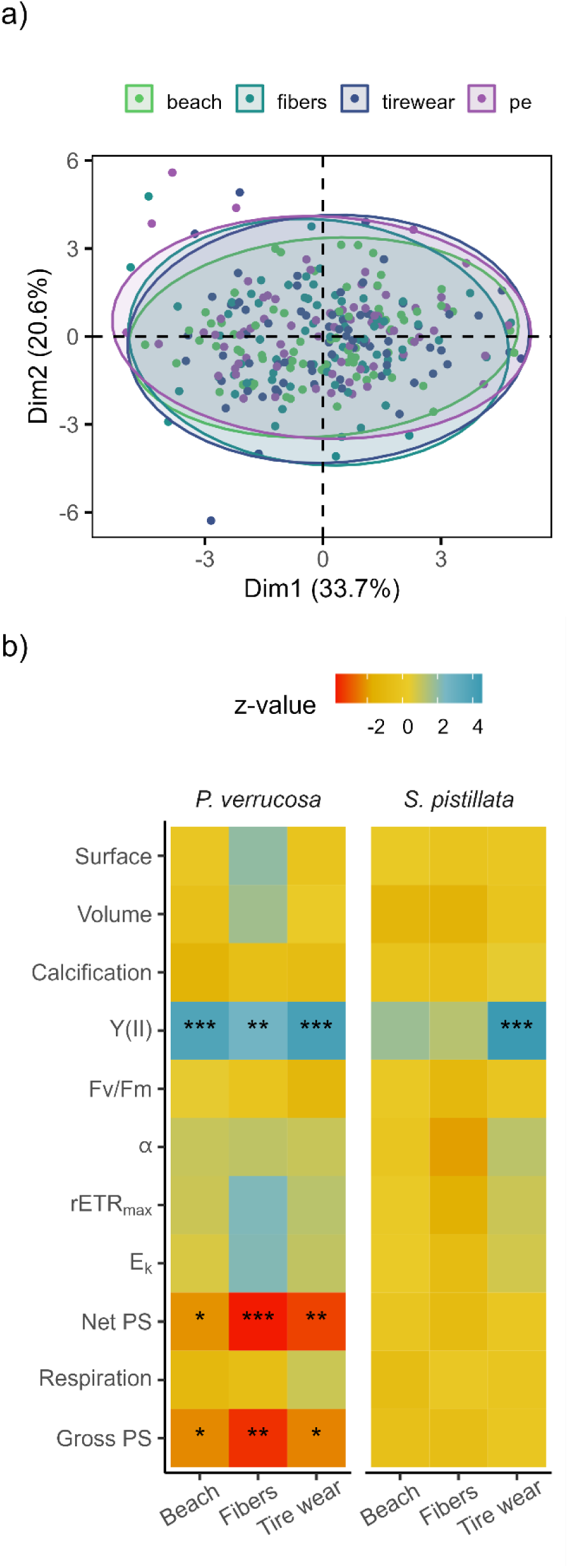
Effects of the single polymer treatment PE in comparison to the three complex microdebris treatments (beach, fibers, and tire wear). a) Principal component analyses (PCA) are constructed using 12 physiological parameters (growth in surface area, growth in volume, calcification, Y(II), Fv/Fm, rETR_max_, E_k_, α, net photosynthesis, respiration, and gross photosynthesis) of *P. verrucosa* and *S. pistillata*. b) Statistical differences (z-values) between the PE treatment and the complex microdebris treatments are illustrated as heatmap with higher values (blue) indicating increasing effects and lower values (red) indicating decreasing effects compared to the PE treatment. Statistically significant differences are marked with an asterisk, defined as p < 0.001 (***), p < 0.01 (**), and p < 0.05 (*) and derived from linear mixed-effects models followed by holm-adjusted posthoc comparison (n = 18).

### 3.5 Feeding responses to different microdebris

Reaction, uptake, and egestion of corals differed between the microdebris (beach, fibers, tire wear, and PE) and the natural food (copepods), (Figure 6, Table S9)*. P. verrucosa* responded significantly more often to PE (13 %), tire wear (11 %), and beach particles (9 %) than to fibers (3 %); (Fisher’s exact test, PE: p < 0.001, tire wear: p < 0.001, beach: p = 0.019). *S. pistillata* responded similarly to all debris types, in ∼5 % of the cases. Overall, the microdebris elicited significantly less frequent responses from *P. verrucosa* and *S. pistillata* compared to their response to natural food (92 % and 79 %, respectively, Fisher’s exact test, p < 0.001). However, both species handled microdebris significantly longer (*P. verrucosa*: 49 ± 237 sec and *S. pistillata*: 72 ± 316 sec) than natural food particles (*P. verrucosa*: 5 ± 6 sec and *S. pistillata*: 5 ± 7 sec; Wilcox test, both p < 0.001).

**Figure 6:**
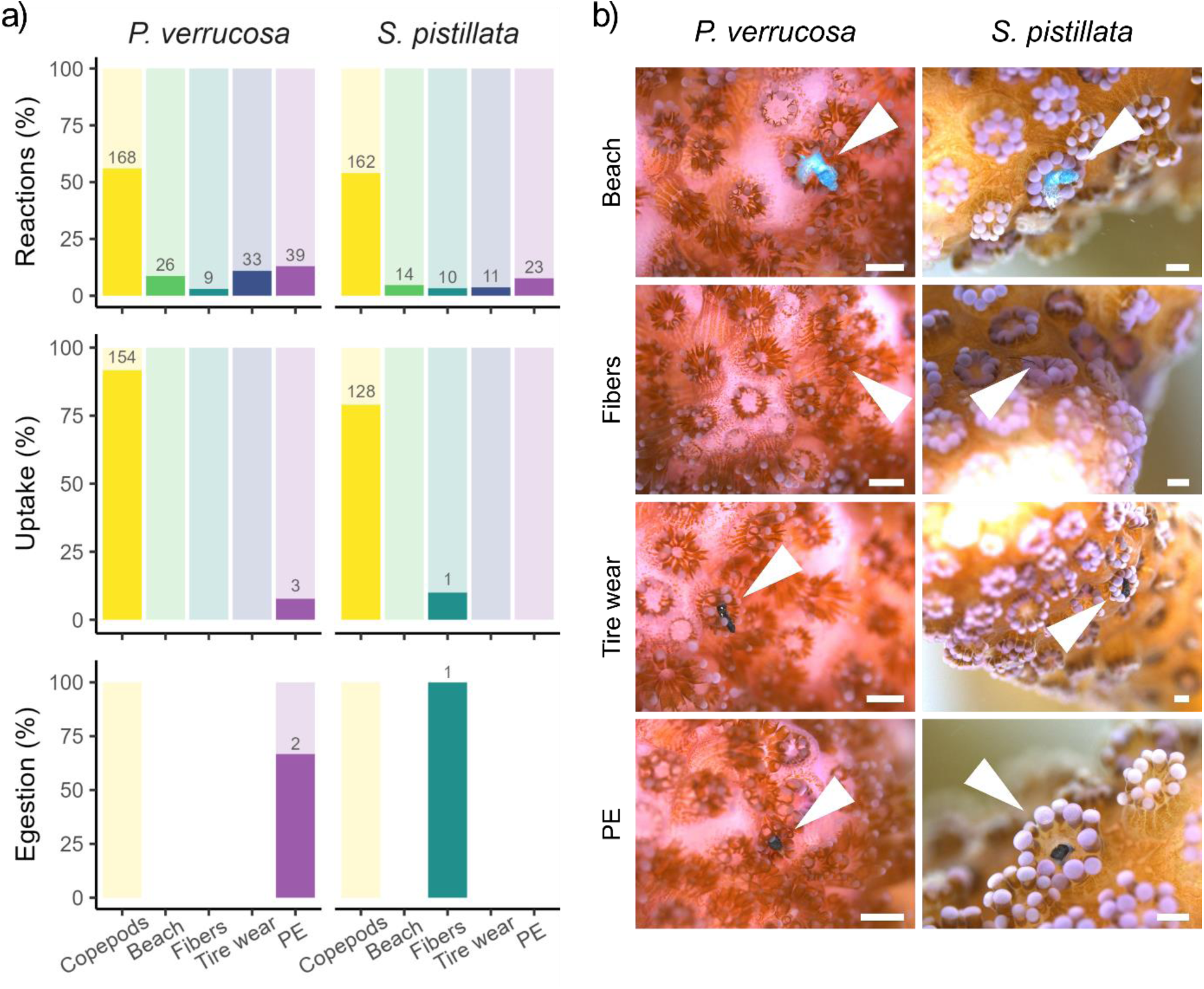
Feeding responses of *P. verrucosa* and *S. pistillata* to natural food (copepods) and microdebris (beach, fibers, tire wear, and PE). a) Frequency of reaction to, uptake, and egestion of copepods and microdebris are shown as bar charts with positive events in dark and negative events in light color, as % of all trials (n = 300 offered particles per species and food type). b) Reactions of both species to the different particles. Arrows indicate the interacting polyp. Scale bars: 500 µm.

Of 300 PE particles offered to *P. verrucosa* three particles were ingested, of which two were egested after 10 sec and 5 min, respectively. *S. pistillata* ingested only one fiber, which was egested after 13 sec. In contrast, most of the copepods were ingested (∼86 %), which was significantly higher than the ingestion of microdebris (Fisher’s exact test, p < 0.001). Natural food particles were generally taken up within 5 ± 2 sec by both species. In contrast, the uptake of microdebris took much longer for *P. verrucosa* (6 ± 1 min, n = 3; Wilcox test, p = 0.026, Table S10). Natural food particles were not released after ingestion in any case.

*P. verrucosa* reacted significantly more often to microdebris than *S. pistillata* (Fisher’s exact test, p < 0.001, Table S11). Specifically, *P. verrucosa* reacted significantly more often to tire wear and PE than *S. pistillata* (Fisher’s exact test, p = 0.001 and p = 0.044, respectively). Both *P. verrucosa* and *S. pistillata* reacted equally often to natural food particles (∼55 %), yet *P. verrucosa* ingested significantly more than *S. pistillata* (92 % vs. 79 % Fisher’s exact test, p > 0.050).

## 4 Discussion

Microdebris occur in complex mixtures in coral reefs, yet experimental approaches to date have focused on the effects of single polymers on stony corals. This is the first study that investigated the effects of complex microdebris mixtures. While most microdebris types had no significant physiological effects, parameter- and species-specific effects occurred in some of the treatments. Overall, the fibers and the tire wear treatment had the strongest effects, and *P. verrucosa* was more affected than *S. pistillata*. Positive effects of the microdebris treatments were observed on volume growth in both species, accompanied by decreased calcification rates in *P. verrucosa*. Both species responded with an upregulation in photosynthesis but showed no signs of compromised health. In comparison, the single polymer treatment PE and the complex microdebris treatments had similar, yet again parameter- and species-specific effects on coral physiology. The two species also differed in their feeding responses to the particles, where *P. verrucosa* reacted significantly more often to the microdebris than *S. pistillata*.

### 4.1 Microdebris treatments have differential effects on coral physiology

In some cases, microdebris caused increased growth in skeletal volume in both species. Specifically, volume growth of *P. verrucosa* was increased in the fibers treatment while calcification decreased in the fibers and beach treatment. This might indicate an overgrowth of particles, as observed in previous studies (Hierl et al., 2021; Reichert et al., 2022, 2018), potentially modifying skeletal structure and leading to increased growth in volume despite decreased calcification. In contrast, photosynthetic rates remained largely unaffected by the exposure to microdebris. Stronger effects were observed in the photosynthetic efficiency of the associated photosymbionts, which was enhanced in both species by the exposure to microdebris. These observations correspond to enhanced photosynthetic efficiency of corals exposed to single polymer particles (Lanctôt et al., 2020; Reichert et al., 2019), but their cause remains a subject of speculation. This upregulation might be a mechanism to compensate for impaired feeding success and associated loss of energy. It appears to successfully mitigate the impacts of microdebris as the exposure had no effects on surface growth and coral health (i.e., necrosis). Another reason for the altered photochemistry might be that the presence of microdebris might have subtle impacts on the light conditions (both intensity and/or spectral composition), which remains to be assessed.

Overall, the microdebris treatments caused only occasional, minor impacts on the physiology of the tested coral species, which were often masked by the large biological variation between colonies. This is in line with previous studies on the effects of microplastics on corals exposed to moderate concentrations over the timeframe of weeks to months (Reichert et al., 2021, 2019). This suggests that the concentration tested here only constitutes a minor, sublethal stressor for corals, which seem to possess mechanisms to cope with the exposure over the tested timeframe. Yet, some specific effects occurred and artificial fibers from clothing, in particular, induced the most changes in coral physiology. Reasons for this might be that fibers frequently entangle on the coral colonies, and cleaning mechanisms that might be effective for microdebris fragments fail to work (Hierl et al., 2021; Oldenburg et al., 2021; Rades et al., 2022; Reichert et al., 2018). Once fibers are entangled, leaching chemicals might be directly transferred to the neighboring tissue (Aminot et al., 2020; Montano et al., 2020; Saliu et al., 2019). If the exposure is prolonged over several months, it might cause necrosis (Hierl et al., 2021; Reichert et al., 2019). This has, however, not been observed during the 8 weeks of exposure in this study. Rather it appeared that fibers, which could not be removed by the corals, were overgrown, causing increased volume growth. The stronger effects of these treatments observed in this study might be partly caused by the higher amount of fibers in the water column (Fig. S1). However, *P. verrucosa* responded significantly less often to fibers than to fragments, which suggests that the negative effects are not caused by the reaction to and the ingestion of fibers, but rather by the entanglement and potentially resulting interferences (i.e., movement impairments of polyps, mechanical disturbance or abrasion, and toxin or pathogen transfer to the coral surface). Tire wear and residues from the automobile sector also caused several effects, specifically on the photosynthetic efficiency of the symbionts. This can hint towards a chemical leaching (Carrasco-Navarro et al., 2022; Jeong et al., 2022), especially as the tire wear particles, which seemed to sink to the ground due to their negative buoyancy, resulted in the lowest concentration of particles in the water column. The major constituents of tire wear are synthetic and natural rubber, and vulcanization agents such as sulfur, zinc, polyaromatic hydrocarbons, and thiazoles (Wagner et al., 2018). Tire wear has been shown to leach considerable amounts of these substances, which are known to be toxic to corals and other organisms (Halle et al., 2020; Ouédraogo et al., 2021) and have been identified to exhibit a higher toxicity than other plastic polymers (Sørensen et al., 2023). In contrast, the effect of the beach treatment appeared rather low, comparable to the impact of the PE treatment. These findings are supported by the observation that the corals responded as often to the beach particles as to all other fragments. Further, this could be explained by the fact that the beach treatment primarily consisted of PE (∼60 %, Kühn et al., 2018) and that the treatment consisted of environmental microdebris that had been collected from the seaside, where their chemical load had likely already reduced over time.

### 4.2 Single polymer treatments are representative of complex microdebris types but differ in specifics

The PE treatment had no consistently different effect compared to the mixed microdebris treatments. Yet parameter- and species-specific differences occurred. In particular, when exposed to PE, *P. verrucosa* showed no upregulation in photosynthetic efficiency as observed for other microdebris treatments, but in photosynthesis. This indicates that corals respond differently to complex microdebris compared to single polymer microdebris. This is likely due to the different physical and chemical properties found in the microdebris treatments (i.e., density, shape, polymer composition, and additives). Although the results from single polymer experiments were not generally different, future laboratory experiments should rather use mixed microdebris to mimic naturally occurring microdebris and assess the effects of microdebris on corals in more detail.

### 4.3 Species-specific effects might be related to particle detection

The effects of the microdebris treatments were species-specific, which is in line with previous observations on single polymer exposure experiments (Mendrik et al., 2021; Mouchi et al., 2019; Reichert et al., 2021, 2019). Overall, the physiology of *S. pistillata* was less strongly affected by the microdebris exposure than *P. verrucosa*. To date, reasons for species-specific effects are poorly understood but hint towards differences in colony morphology and feeding strategy of the coral species (Hankins et al., 2022; Martin et al., 2019; Rades et al., 2022; Reichert et al., 2022). Here we find that particle detection and handling might be a reason for the observed differences. The interaction might hinder feeding and be energetically costly. This might also explain why the upregulation in photosynthetic efficiency is more pronounced (i.e., affected more parameters) in *P. verrucosa*, although it occurred earlier (after 4 weeks) in *S. pistillata*. The lower reaction rates of *S. pistillata* to microdebris than of *P. verrucosa* suggest differences in particle detection mechanisms between the species. These may underlie the differences observed here, with better compensation strategies to environmental impacts in *S. pistillata*, as previously observed with other stressors, such as thermal stress and bleaching (Kvitt et al., 2011; Meziere et al., 2022).

#### Implications for coral reef conservation

Taken together, we found that microdebris had minor impacts on reef-building corals in our study, yet, the fibers and tire wear treatments impacted multiple physiological parameters. Although these effects had no detrimental impacts on coral health, they might add up to the already existing stress posed on coral reef ecosystems and may push coral reef communities past their tipping points (Mendrik et al., 2021; Reichert et al., 2021). This highlights the urgency for stronger regulations of microdebris influxes from land (Trudsø et al., 2022). While methods to effectively remove these residues from our wastewater are already available, they need to be implemented more broadly to protect coral reefs from the effects of microdebris. Thus, we call for prioritized measures to reduce residues from artificial clothes and the automobile sector. This should be considered in both our individual actions as well as policy-making.

### 4.4 Conclusions

Our study is the first to present the effects of common microdebris types from major sources of anthropogenic pollution on two reef-building coral species. We show that residues from artificial clothes and the automobile sector had the strongest effects on the tested corals and that the effects of these mixtures might be more pronounced than those of single polymers. Although single polymer treatments appeared overall representative to investigate the effects of microdebris pollution, the underlying physiological mechanisms of how corals respond to naturally occurring complex microdebris might not be resolved, as different mixtures seem to cause different responses. Reasons for species-specific differences in the effects of microdebris on corals might be attributed to different feeding strategies and the capability to reject unwanted particles. However, this warrants further evidence. Our study underlines the urgency to implement effective wastewater treatments and address the major sources of pollution for microdebris.

## 6 Acknowledgments

We thank Birgit Zettl and Christina Anding from Justus Liebig University Giessen, Germany for the animal care and maintenance. The study was conducted as part of the ‘Ocean 2100’ global change simulation project of the Colombian-German Center of Excellence in Marine Sciences (CEMarin). The ‘beach’ treatment material was prepared within the JPI Oceans ‘PLASTOX’ project.

## 7 Author Contributions

JR: Conceptualization, Formal analysis, Investigation, Writing – original draft, review & editing. VT: Conceptualization, Formal analysis; Investigation, Writing – review & editing. KP, EW, LB, SP: Investigation. SK: Resources, Writing – review & editing. PS, MZ, TW: Conceptualization, Resources, Writing – review & editing.

## 8 Data and Materials Availability

The analyzed data and 3D models of the studied corals are available at figshare.com: https://doi.org/10.6084/m9.figshare.23983125.v1.

## 9 Use of generative AI

During the preparation of the graphical abstract, the authors used Canva Text-to-image in order to generate an illustration of a coral reef. After using this tool, the authors reviewed the content as needed and take full responsibility for the content of the publication.

