## Supplementary Materials for "Common types of microdebris affect the physiology of reef-building corals"

### Affiliation

S1 Supplementary material and methods

S1.1 Detailed description of experimental setup

Coral colonies were maintained at the 'Ocean2100' coral facility at Justus Liebig University Giessen, Germany, under laboratory conditions (in accordance with the institutional animals' care guidelines, long-term rearing: 10:14 light:dark photoperiod, light intensity (PAR) 200 µmol photons $m^{-2}$ $s^{-1}$, and temperature 26 ± 0.5 °C) for at least six months before the experiment (see details on coral colonies in Table S1). Twelve coral fragments were placed in each tank for this study together with additional 6 fragments that were part of a different study (Reichert et al., 2021).

A wavemaker pump with a maximum water turnover of 2800-4000 $L \cdot h^{-1}$ (easySTREAM pro Wavemaker Pump, ES-18, Aqualight GmbH, Germany) at mode 4 and a power of 10/20 together with a turnover pump with a maximum water turnover of 700 $L \cdot h^{-1}$ (Submarine Water Pump, Resun S-700, Resun, China) generated steady water movement and distribution. The average velocimetry at the corals' position was 3.8 $m \cdot s^{-1}$. To submerge microdebris particles accumulating at the water surface, the outflow piece of the turnover pump was extended by a u-shaped PVC tube, which ended approximately 1 cm above the water surface. Water temperature was feedback-controlled in each tank using sensors (GHL Temp Sensor digital, GHL Advanced Technology GmbH & Co. KG, Germany) connected to GHL aquarium computers (ProfiLux 3 and ProfiLux 4, GHL Advanced Technology GmbH & Co. KG, Germany) and a submersible titanium heater (Schego Heater 300 W, Protection class IP 68, SCHEGO Schemel & Goetz GmbH & Co KG, Germany). Water temperatures were maintained at 27 ± 0.3 °C.

Above each tank four LED lamps with different combinations of white and blue LED spots were installed ~32 cm above the water surface (SunaECO LED Lighting Strip: Marine Blue, Marine White, Reef Blue, Reef White, AquaRay by Tropical Marine Centre, United Kingdom).

Light was provided with an average PAR intensity of 120 µmol photons m$^{-2}$·s$^{-1}$ for 10/24 hours (from 8 am to 6 pm). The left and right outer sides of the tanks were masked with a dark blue adhesive foil to minimize light dispersion.

Two self-made outflow filters with a mesh size of 65 µm were installed in each tank to prevent microdebris particles from leaving and contaminating other aquaria. The experimental tanks were connected to a technical tank (265 L) and eight other tanks of different sizes building one system with a total water volume of about 2,300 L. Water coming from the experimental tanks was filtered through a fleece filter bag as a second particle removal safety measure. To remove organic refuse, the system was equipped with a protein skimmer (ATI PowerCone 250i, ATI-Aquaristik, Germany). For the adsorption of phosphate, a silicate filter (ATI-Aquaristik, Germany) was filled with iron oxide granulate (AquaLight PHOS, 0.5–2 mm, Aqualight GmbH, Germany). To perpetuate a constant level of calcium concentration and alkalinity a calcium reactor was connected to the technical tank. The flow-through of the reactor was regulated using feedback-controlled alkalinity measurements (six times in 24 h) through an Alkatronic (Alkatronic – Alkalinity, Whitecorals Vertriebs-GmbH, Germany). A UVC-sterilizer (RWUVC/20/1000, RuWal aquatech, Italy) was added to reduce parasites and pathogens in the whole system with a turnover rate of 1,000 L·h$^{-1}$ using ultraviolet radiation (intensity: 33,000 µm cm$^{-2}$·s$^{1}$). In a separate compartment of the technical tank, an algal refugium was arranged using *Chaetomorpha* sp. to buffer pH levels overnight and reduce algae growth by removing excess nutrients from the water. The light in the refugium had a contrariwise cycle to the light of the experimental aquaria.

Water inflow was maintained at approximately 78 L·h$^{-1}$ using an inflow pump (easyPump 24V-DC, EP-12000, 3,600-12,000 L·h$^{-1}$, Aqualight GmbH, Germany). Incoming water was filtered before entering the tanks through a mesh (mesh size: 65 µm) to minimize contamination through algae or detritus from the connected system. Evaporated water was replaced automatically in the technical tank six times a day.

S1.2 Maintenance of the experiment

Tanks were checked daily to maintain the experiment. In- and outflow filters were cleaned with hot water at least once a day. Before filters were cleaned, particles stuck on the sides were rinsed off with saltwater to keep them in the tanks. In case of strong algae accumulation on the filters, they were placed in NaOCl (Danklorix, CP GABA GmbH, Germany; dilution 0.5 L with 15 L water) for 20 minutes to remove strongly adhering organic material. The corals were fed daily with copepods (Calanoide Copepoden, Zooschatz, Germany). For this, approximately 0.53 g of copepods (frozen feed) were given to each tank. To provide additional amino acids, around 0.53 ml amino acid mixture (Pohl's Xtra special, Korallenzucht.de Vertriebs GmbH, Germany) was added daily to the water column. Whenever necessary (i.e., visible algae growth and before measurements) fishing lines of the corals, were cleaned from algae. To ensure constant water conditions, water parameters (i.e., salinity, temperature, carbonate hardness, phosphate, calcium, nitrate, nitrite, and magnesium), were measured regularly. The salinity was adjusted to 35-36 ‰ and controlled daily with a refractometer (HI 96822 Seawater Refractometer, Hanna Instruments Ltd, United Kingdom). Carbonate hardness was maintained at 1.424 $mmol \cdot L^{-1}$ (8 °dKH) tested and controlled via an Alkatronic (Alkatronic – Alkalinity, Whitecorals Vertriebs-GmbH, Germany). Phosphate ($PO_4^{3-}$) was kept below 0.02 $mg \cdot L^{-1}$, measured with a Spectroquant Phosphate Test (10 ml sample, Merck KGaA, Germany). Calcium had an average concentration of 400 $mg \cdot L^{-1}$. It was measured once a week using titration with EDTA. Nitrate ($NO_3^-$) and nitrite ($NO_2^-$) were below detectable ranges and were checked weekly using Nitrate Test Merckoquant 110020 (Merck KGaA, Germany). Magnesium concentration had an average of 1,300 $mg \cdot L^{-1}$ and was measured every 4 weeks with a titration test (Mg Profi Test, Salifert, The Netherlands).

S2 Supplementary figures

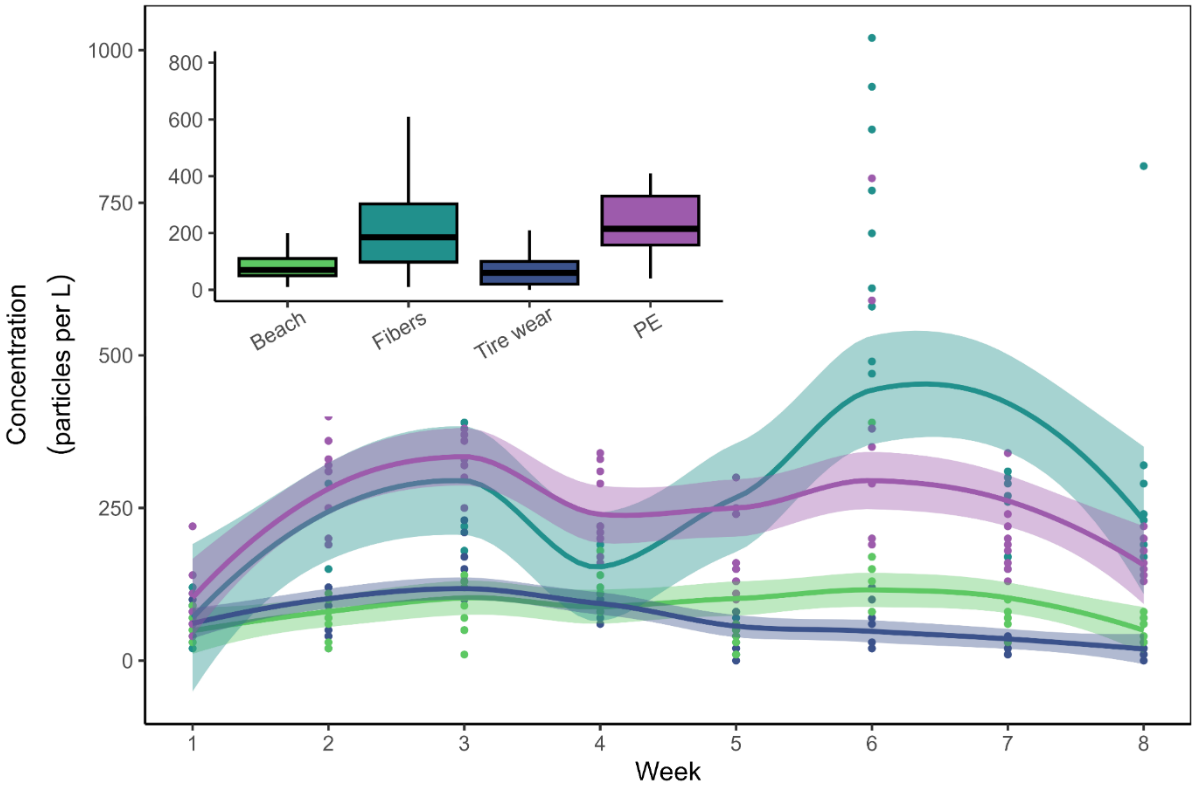


Figure S1: Microdebris particle concentrations (particles per L). Development of particle concentrations is shown as raw data points and local polynomial regression fitting (solid line) with 95 % confidence intervals (colored area). Boxplots display average concentrations from all time points.

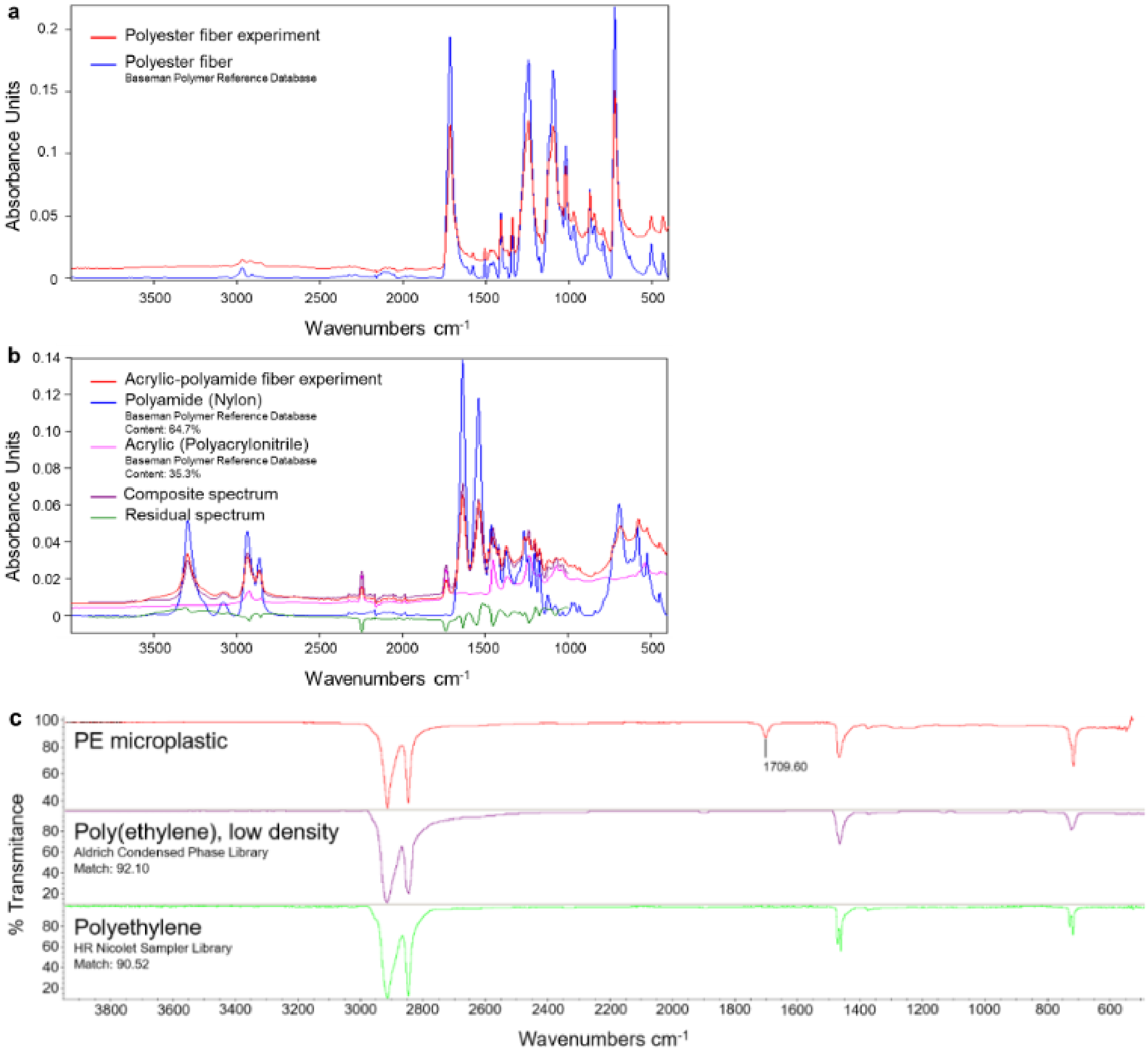


Figure S2: FTIR spectra of fibers (a: polyester and b: acrylic-polyamide) and PE microplastic particles (c) used in the experiment (red) are compared to reference spectra (other colors).

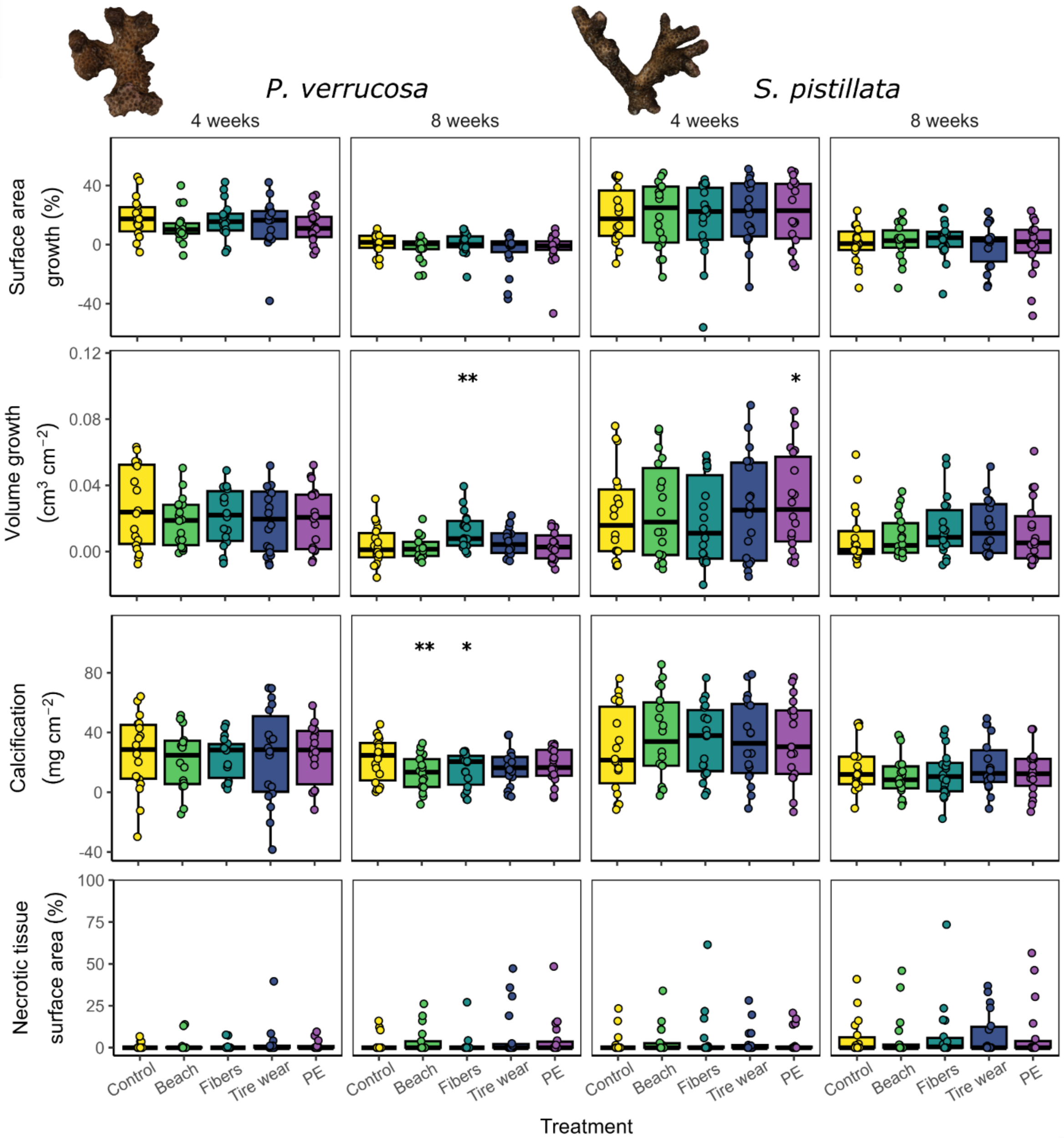


Figure S3: Growth rates (surface area, volume, and calcification) and necrosis of *P. verrucosa* and *S. pistillata* exposed to the different treatments (control, beach, fibers, tire wear, and PE) are shown as boxplots. Rates are shown for the first 4 and the following 4 weeks of the experiment. Data are displayed as box-and-whisker plots with raw data points; lines indicate medians, boxes indicate the first and third quartile, and whiskers indicate ± 1.5 IQR. Significant differences between the control and the microdebris treatments are derived from linear mixed-effects models (surface area growth, volume growth, calcification) or generalized linear mixed-effects models (necrotic tissue surface area) followed by holm-adjusted posthoc comparison ($n = 18$) and are defined as $p < 0.001$ (***), $p < 0.01$ (**), and $p < 0.05$ (*).

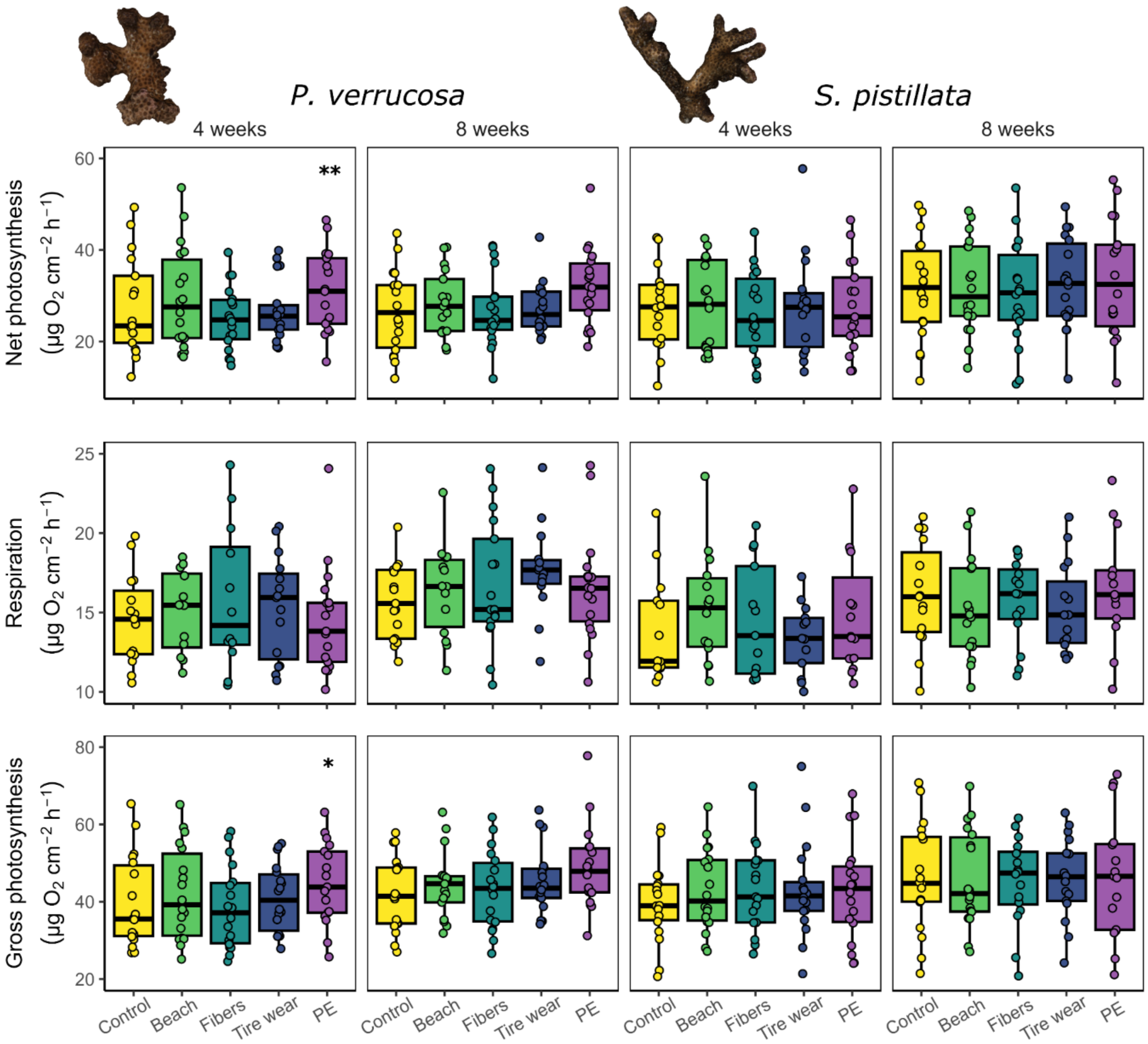


Figure S4: Photosynthesis rates (net photosynthesis, respiration, and gross photosynthesis) of *P. verrucosa* and *S. pistillata* exposed to the different treatments (control, beach, fibers, tire wear, and PE) are shown as boxplots. Rates are shown for the first 4 and the following 4 weeks of the experiment. Data are displayed as box-and-whisker plots with raw data points; lines indicate medians, boxes indicate the first and third quartile, and whiskers indicate ± 1.5 IQR. Significant differences between the control and the microdebris treatments are derived from linear mixed-effects models followed by holm-adjusted posthoc comparison ($n = 18$) and are defined as $p < 0.001$ (***), $p < 0.01$ (**), and $p < 0.05$ (*).

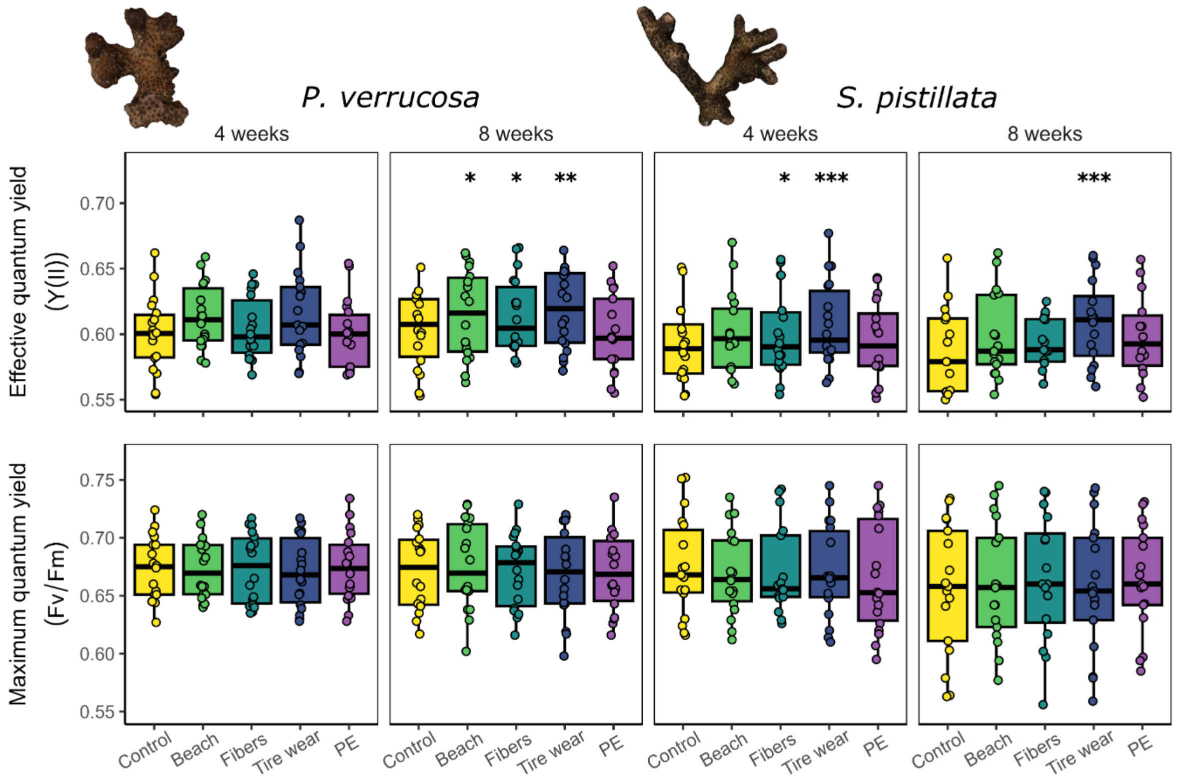


Figure S5: Photosynthetic efficiency (effective and maximum quantum yield) of *P. verrucosa* and *S. pistillata* exposed to the different treatments (control, beach, fibers, tire wear, and PE) are shown as boxplots. Rates are shown for the first 4 and the following 4 weeks of the experiment. Data are displayed as box-and-whisker plots with raw data points; lines indicate medians, boxes indicate the first and third quartile, and whiskers indicate ± 1.5 IQR. Significant differences between the control and the microdebris treatments are derived from linear mixed-effects models followed by holm-adjusted posthoc comparison (n = 18) and are defined as $p < 0.001$ (***), $p < 0.01$ (**), and $p < 0.05$ (*).

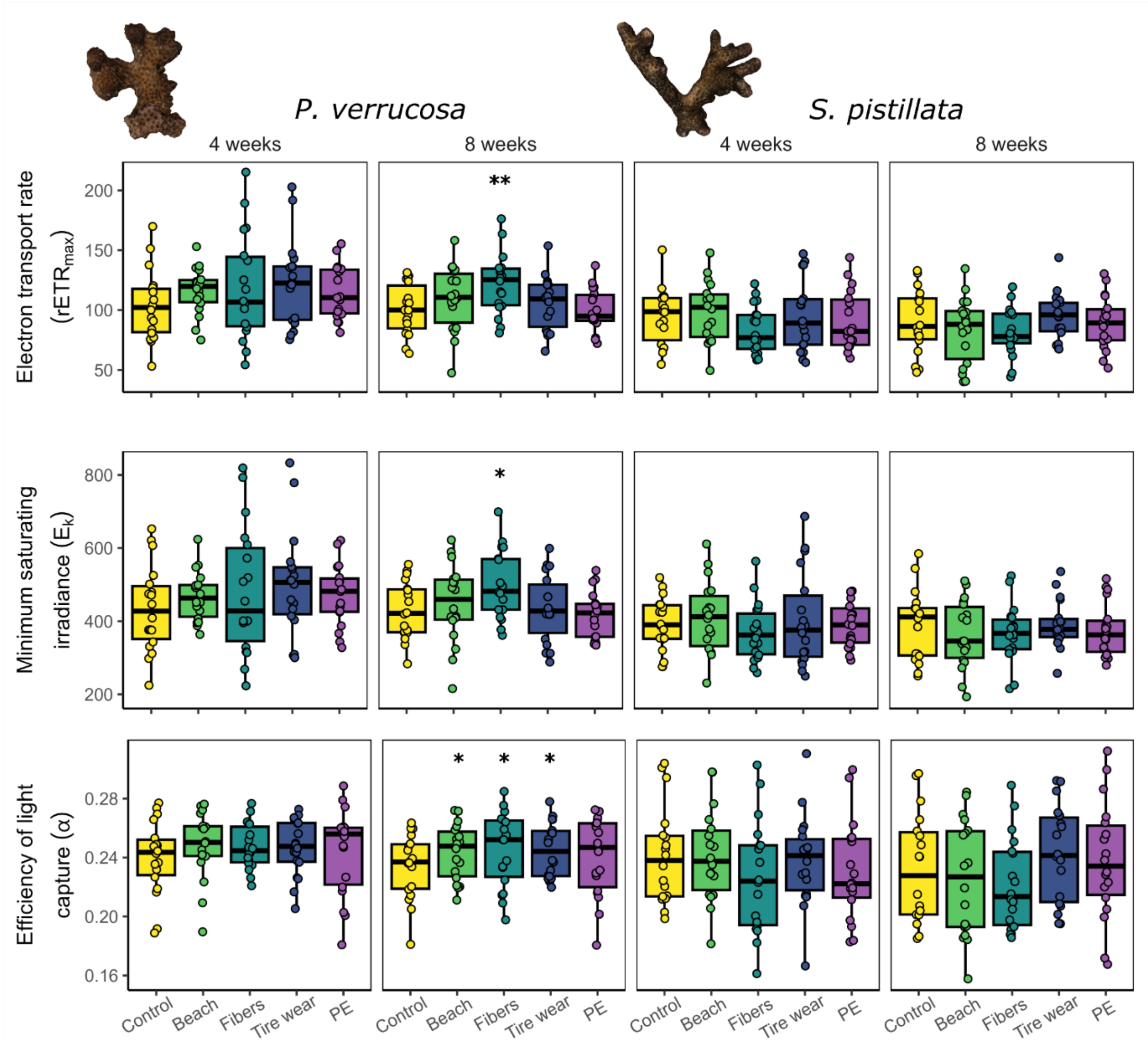


Figure S6: Photosynthetic efficiency derived from rapid light curves (electron transport rate, minimum saturation irradiance, and efficiency of light capture) of *P. verrucosa* and *S. pistillata* exposed to the different treatments (control, beach, fibers, tire wear, and PE) are shown as boxplots. Rates are shown for the first 4 and the following 4 weeks of the experiment. Data are displayed as box-and-whisker plots with raw data points; lines indicate medians, boxes indicate the first and third quartile, and whiskers indicate ± 1.5 IQR. Significant differences between the control and the microdebris treatments are derived from linear mixed-effects models followed by holm-adjusted posthoc comparison ($n = 18$) and are defined as $p < 0.001$ (***), $p < 0.01$ (**), and $p < 0.05$ (*).

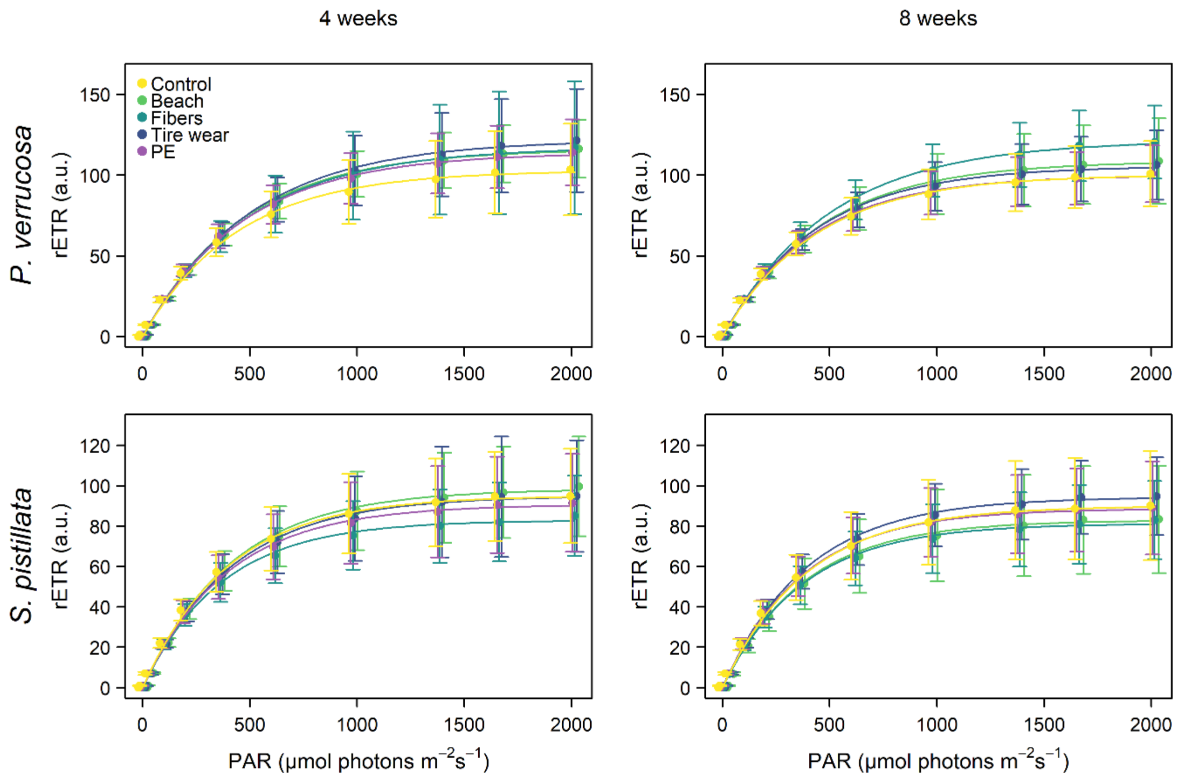


Figure S7: Rapid light curves of *P. verrucosa* and *S. pistillata* exposed to different microdebris treatments (control, beach, fibers, tire wear, PE). Values of measured relative electron transport rate (rETR) for the used PAR intensities were measured after 4 and 8 weeks.

S3 Supplementary tables

Table S1: Origin of the studied coral colonies, month of collection, and entering the facility together with the CITES registration numbers.

| Species | Colony | Origin | Collection | Entering | CITES number |
|---|---|---|---|---|---|
| *P. verrucosa* | A | Indonesia | 12/2007 | 12/2007 | 14846/IV/SATS-LN/2007 |
| *P. verrucosa* | B | Indonesia | 04/2014 | 05/2014 | 14NL214371/11 |
| *P. verrucosa* | C | Indonesia | 12/2007 | 12/2007 | 14846/IV/SATS-LN/2007 |
| *P. verrucosa* | D | Saudi Arabia | 03/2019 | 03/2019 | 19-SA-000091-PD |
| *P. verrucosa* | E | Saudi Arabia | 03/2019 | 03/2019 | 19-SA-000091-PD |
| *P. verrucosa* | F | Saudi Arabia | 03/2019 | 03/2019 | 19-SA-000091-PD |
| *S. pistillata* | A | | > 10 years | > 10 years | |
| *S. pistillata* | B | | > 10 years | > 10 years | |
| *S. pistillata* | C | Fiji | 2016 | 2016 | 15NL229192/11 |
| *S. pistillata* | D | Saudi Arabia | 03/2019 | 03/2019 | 19-SA-000092-PD |
| *S. pistillata* | E | Saudi Arabia | 03/2019 | 03/2019 | 19-SA-000092-PD |
| *S. pistillata* | F | Saudi Arabia | 03/2019 | 03/2019 | 19-SA-000092-PD |

Table S2: Settings used for the calculation and documentation of 3D models of the corals in Artec Studio 11.

| Process | Step | Setting |
|---|---|---|
| Fine registration | Registration algorithm | Texture and Geometry |
| | Refine serial | On |
| | Loop closure | Off |
| Global registration | Registration algorithm | Texture and Geometry |
| | Minimal distance | 2 |
| | Iterations | 30 000 |
| Outlier removal | Std dev ml threshold | 3 |
| | Resolution | 0.2 |
| Sharp fusion | Resolution | 0.2 |
| | Fill holes | Watertight |
| | Remove targets | Off |
| Texture | Generate triangle map | On |
| | Triangle size (in pixels) | 10 |
| | Enable texture normalization | Off |

Table S3: PAM settings used for all measurements (relative effective yield, relative maximum yield, rapid light curves) for duration and intensity of measuring light, as well as saturation pulse.

| Category | Parameter | Setting |
|---|---|---|
| Measuring Light | Intensity | 4 |
| | MF-L. | 200 |
| | MF-H. | 20,000 |
| Actinic Light | Intensity | 1 |
| | Width (s) | 0 |
| | Color | Red |
| PS I Light | Intensity | 4 |
| | Width (s) | 5 |
| | Color | Far Red |
| Saturation Pulse / MT | Intensity | 8 |
| | Intensity (Fo, Fm) | 6 |
| | SP-Width (10 ms) | 8 |
| Slow Induction | Delay (s) | 40 |
| | Clock (s) | 20 |
| | Width (s) | 310 |
| Analysis Mode | | SP-Analysis |

Table S4: Results of statistical analyses assessing the effects of the microdebris treatments (beach, fibers, tire wear, and PE) compared to the control on growth (i.e., surface area, volume, and calcification) and health parameters (i.e., necrosis) of *Pocillopora verrucosa* and *Stylophora pistillata* in the first 4 weeks of the experiment (t0–t1) and in the following 4 weeks of the experiment (t1–t2). Results are derived from (generalized) linear mixed effects models with treatment as a fixed factor and colony (col) set as a random factor, on scale- or log-transformed data, followed by holm-adjustment for multiple testing. Numbers of observations (n), estimates, standard errors, z-values, and p-values are given with model specifications. Bold values indicate significance ($p < 0.05$).

| Species | Response variable | Time | Contrast | | | Estimate | Std. Error | z-value | p | n | Model specifications |
|---|---|---|---|---|---|---|---|---|---|---|---|
| *P. verrucosa* | Surface area | t0-t1 | control | - | beach | 0.444 | 0.226 | 1.964 | 0.198 | 89 | lmer(scale(x)~treatment + (1\|col)) |
| | | | control | - | fibers | 0.097 | 0.226 | 0.429 | 0.668 | | |
| | | | control | - | tire wear | 0.379 | 0.226 | 1.674 | 0.198 | | |
| | | | control | - | PE | 0.426 | 0.226 | 1.883 | 0.198 | | |
| | | t1-t2 | control | - | beach | 0.329 | 0.241 | 1.361 | 0.389 | 88 | lmer(scale(x)~treatment + (1\|col)) |
| | | | control | - | fibers | 0.013 | 0.238 | 0.055 | 0.956 | | |
| | | | control | - | tire wear | 0.504 | 0.238 | 2.121 | 0.136 | | |
| | | | control | - | PE | 0.360 | 0.238 | 1.516 | 0.389 | | |
| *S. pistillata* | Surface area | t0-t1 | control | - | beach | -0.021 | 0.158 | -0.135 | 1.000 | 90 | lmer(scale(x)~treatment + (1\|col)) |
| | | | control | - | fibers | 0.144 | 0.158 | 0.912 | 1.000 | | |
| | | | control | - | tire wear | -0.091 | 0.158 | -0.575 | 1.000 | | |
| | | | control | - | PE | -0.101 | 0.158 | -0.637 | 1.000 | | |
| | | t1-t2 | control | - | beach | -0.115 | 0.202 | -0.566 | 1.000 | 90 | lmer(scale(x)~treatment + (1\|col)) |
| | | | control | - | fibers | -0.226 | 0.202 | -1.117 | 1.000 | | |
| | | | control | - | tire wear | 0.056 | 0.202 | 0.276 | 1.000 | | |
| | | | control | - | PE | 0.069 | 0.202 | 0.343 | 1.000 | | |
| *P. verrucosa* | Volume | t0-t1 | control | - | beach | 0.472 | 0.208 | 2.269 | 0.093 | 89 | lmer(scale(x)~treatment + (1\|col)) |
| | | | control | - | fibers | 0.333 | 0.208 | 1.601 | 0.169 | | |
| | | | control | - | tire wear | 0.425 | 0.208 | 2.044 | 0.123 | | |
| | | | control | - | PE | 0.359 | 0.208 | 1.724 | 0.169 | | |
| | | t1-t2 | control | - | beach | 0.086 | 0.275 | 0.314 | 1.000 | 88 | lmer(scale(x)~treatment + (1\|col)) |
| | | | control | - | fibers | -0.855 | 0.270 | -3.161 | **0.006** | | |

| | | | | | | | | | | | |
|---|---|---|---|---|---|---|---|---|---|---|---|
| | | | control | - | tire wear | -0.217 | 0.270 | -0.803 | 1.000 | | |
| | | | control | - | PE | 0.037 | 0.270 | 0.136 | 1.000 | | |
| *S. pistillata* | Volume | t0-t1 | control | - | beach | -0.116 | 0.117 | -0.994 | 0.636 | 90 | lmer(scale(x)~treatment + (1\|col)) |
| | | | control | - | fibers | 0.117 | 0.117 | 0.999 | 0.636 | | |
| | | | control | - | tire wear | -0.150 | 0.117 | -1.285 | 0.596 | | |
| | | | control | - | PE | -0.293 | 0.117 | -2.502 | **0.049** | | |
| | | t1-t2 | control | - | beach | 0.007 | 0.203 | 0.034 | 0.973 | 90 | lmer(scale(x)~treatment + (1\|col)) |
| | | | control | - | fibers | -0.318 | 0.203 | -1.569 | 0.441 | | |
| | | | control | - | tire wear | -0.323 | 0.203 | -1.597 | 0.441 | | |
| | | | control | - | PE | -0.166 | 0.203 | -0.819 | 0.825 | | |
| *P. verrucosa* | Calcification | t0-t1 | control | - | beach | 0.153 | 0.202 | 0.756 | 1.000 | 90 | lmer(scale(x)~treatment + (1\|col)) |
| | | | control | - | fibers | 0.070 | 0.202 | 0.346 | 1.000 | | |
| | | | control | - | tire wear | 0.034 | 0.202 | 0.167 | 1.000 | | |
| | | | control | - | PE | 0.005 | 0.202 | 0.026 | 1.000 | | |
| | | t1-t2 | control | - | beach | 0.684 | 0.207 | 3.303 | **0.004** | 89 | lmer(scale(x)~treatment + (1\|col)) |
| | | | control | - | fibers | 0.546 | 0.207 | 2.638 | **0.025** | | |
| | | | control | - | tire wear | 0.430 | 0.207 | 2.075 | 0.076 | | |
| | | | control | - | PE | 0.336 | 0.207 | 1.623 | 0.105 | | |
| *S. pistillata* | Calcification | t0-t1 | control | - | beach | -0.330 | 0.141 | -2.342 | 0.077 | 89 | lmer(scale((x))~treatment + (1\|col)) |
| | | | control | - | fibers | -0.233 | 0.141 | -1.652 | 0.177 | | |
| | | | control | - | tire wear | -0.240 | 0.141 | -1.704 | 0.177 | | |
| | | | control | - | PE | -0.323 | 0.143 | -2.261 | 0.077 | | |
| | | t1-t2 | control | - | beach | 0.388 | 0.170 | 2.281 | 0.090 | 89 | lmer(scale(x)~treatment + (1\|col)) |
| | | | control | - | fibers | 0.317 | 0.170 | 1.861 | 0.188 | | |
| | | | control | - | tire wear | -0.030 | 0.170 | -0.174 | 0.862 | | |
| | | | control | - | PE | 0.241 | 0.170 | 1.414 | 0.315 | | |
| *P. verrucosa* | Necrosis | t0-t1 | control | - | beach | -0.008 | 0.033 | -0.229 | 1.000 | 89 | glmer(x+100)~treatment + (1\|col),family="poisson") |
| | | | control | - | fibers | 0.000 | 0.033 | -0.013 | 1.000 | | |
| | | | control | - | tire wear | -0.022 | 0.033 | -0.674 | 1.000 | | |
| | | | control | - | PE | -0.005 | 0.033 | -0.143 | 1.000 | | |

<table>
<tr><th></th><th></th><th></th><th></th><th></th><th></th><th></th><th></th><th></th><th></th><th></th><th></th></tr>
<tr><td></td><td></td><td>t1-t2</td><td>control</td><td>-</td><td>beach</td><td>-0.019</td><td>0.033</td><td>-0.556</td><td>1.000</td><td>89</td><td>glmer(x+100)~treatment + (1|col),family="poisson")</td></tr>
<tr><td></td><td></td><td></td><td>control</td><td>-</td><td>fibers</td><td>0.004</td><td>0.033</td><td>0.119</td><td>1.000</td><td></td><td></td></tr>
<tr><td></td><td></td><td></td><td>control</td><td>-</td><td>tire wear</td><td>-0.052</td><td>0.033</td><td>-1.596</td><td>0.442</td><td></td><td></td></tr>
<tr><td></td><td></td><td></td><td>control</td><td>-</td><td>PE</td><td>-0.030</td><td>0.033</td><td>-0.930</td><td>1.000</td><td></td><td></td></tr>
<tr><td>S. pistillata</td><td>Necrosis</td><td>t0-t1</td><td>control</td><td>-</td><td>beach</td><td>-0.017</td><td>0.033</td><td>-0.511</td><td>1.000</td><td>90</td><td>glmer(x+100)~treatment + (1|col),family="poisson")</td></tr>
<tr><td></td><td></td><td></td><td>control</td><td>-</td><td>fibers</td><td>-0.032</td><td>0.033</td><td>-0.987</td><td>1.000</td><td></td><td></td></tr>
<tr><td></td><td></td><td></td><td>control</td><td>-</td><td>tire wear</td><td>-0.012</td><td>0.033</td><td>-0.357</td><td>1.000</td><td></td><td></td></tr>
<tr><td></td><td></td><td></td><td>control</td><td>-</td><td>PE</td><td>-0.011</td><td>0.033</td><td>-0.329</td><td>1.000</td><td></td><td></td></tr>
<tr><td></td><td></td><td>t1-t2</td><td>control</td><td>-</td><td>beach</td><td>-0.001</td><td>0.032</td><td>-0.040</td><td>1.000</td><td>90</td><td>glmer(x+100)~treatment + (1|col),family="poisson")</td></tr>
<tr><td></td><td></td><td></td><td>control</td><td>-</td><td>fibers</td><td>-0.019</td><td>0.032</td><td>-0.582</td><td>1.000</td><td></td><td></td></tr>
<tr><td></td><td></td><td></td><td>control</td><td>-</td><td>tire wear</td><td>-0.020</td><td>0.032</td><td>-0.614</td><td>1.000</td><td></td><td></td></tr>
<tr><td></td><td></td><td></td><td>control</td><td>-</td><td>PE</td><td>-0.023</td><td>0.032</td><td>-0.722</td><td>1.000</td><td></td><td></td></tr>
</table>

Table S5: Results of statistical analyses assessing the effects of the microdebris treatments (beach, fibers, tire wear, and PE) compared to the control on net photosynthesis, gross photosynthesis, and respiration of *Pocillopora verrucosa* and *Stylophora pistillata* after the first 4 weeks of the experiment (t1) and after 8 weeks of the experiment (t2). Results are derived from linear mixed effects models with treatment as a fixed factor and colony (col) set aa s random factor, on scale- or log-transformed data, followed by holm-adjustment for multiple testing. Numbers of observations (n), estimates, standard errors, z-values, and p-values are given with model specifications. Bold values indicate significance ($p < 0.05$).

| Species | Response variable | Time | Contrast | Estimate | Std. Error | z-value | p | n | Model specifications |
|---|---|---|---|---|---|---|---|---|---|
| *P. verrucosa* | Net photosynthesis | t1 | control - beach | -0.287 | 0.194 | -1.483 | 0.414 | 90 | lmer(scale(x)~treatment + (1\|col) |
| | | | control - fibers | 0.148 | 0.194 | 0.765 | 0.888 | | |
| | | | control - tire wear | -0.014 | 0.194 | -0.075 | 0.940 | | |
| | | | control - PE | -0.620 | 0.194 | -3.204 | **0.005** | | |
| | | t2 | control - beach | -0.084 | 0.088 | -0.950 | 1.000 | 89 | lmer(log(x)~treatment + (1\|col) |
| | | | control - fibers | -0.028 | 0.087 | -0.321 | 1.000 | | |
| | | | control - tire wear | -0.065 | 0.087 | -0.745 | 1.000 | | |
| | | | control - PE | -0.215 | 0.087 | -2.481 | 0.052 | | |
| *S. pistillata* | Net photosynthesis | t1 | control - beach | -0.250 | 0.230 | -1.086 | 1.000 | 90 | lmer(scale(x)~treatment + (1\|col) |
| | | | control - fibers | -0.085 | 0.230 | -0.368 | 1.000 | | |
| | | | control - tire wear | -0.229 | 0.230 | -0.996 | 1.000 | | |
| | | | control - PE | -0.129 | 0.230 | -0.560 | 1.000 | | |
| | | t2 | control - beach | -0.038 | 0.264 | -0.142 | 1.000 | 90 | lmer(scale(x)~treatment + (1\|col) |
| | | | control - fibers | 0.093 | 0.264 | 0.353 | 1.000 | | |
| | | | control - tire wear | -0.102 | 0.264 | -0.387 | 1.000 | | |
| | | | control - PE | -0.194 | 0.264 | -0.735 | 1.000 | | |
| *P. verrucosa* | Respiration | t1 | control - beach | 0.127 | 0.262 | 0.484 | 1.000 | 90 | lmer(scale(x)~treatment + (1\|col) |
| | | | control - fibers | -0.007 | 0.262 | -0.027 | 1.000 | | |
| | | | control - tire wear | -0.208 | 0.262 | -0.792 | 1.000 | | |
| | | | control - PE | -0.174 | 0.262 | -0.664 | 1.000 | | |
| | | t2 | control - beach | -0.170 | 0.313 | -0.543 | 1.000 | 89 | lmer(scale(x)~treatment + (1\|col) |
| | | | control - fibers | -0.183 | 0.308 | -0.593 | 1.000 | | |
| | | | control - tire wear | -0.651 | 0.308 | -2.111 | 0.139 | | |

| | | | | | | | | | |
|---|---|---|---|---|---|---|---|---|---|
| | | | control - PE | -0.287 | 0.308 | -0.929 | 1.000 | | |
| *S. pistillata* | Respiration | t1 | control - beach | -0.054 | 0.094 | -0.575 | 1.000 | 90 | lmer(scale(x)~treatment + (1\|col) |
| | | | control - fibers | -0.073 | 0.094 | -0.784 | 1.000 | | |
| | | | control - tire wear | -0.047 | 0.094 | -0.505 | 1.000 | | |
| | | | control - PE | -0.055 | 0.094 | -0.591 | 1.000 | | |
| | | t2 | control - beach | 0.108 | 0.317 | 0.340 | 1.000 | 90 | lmer(scale(x)~treatment + (1\|col) |
| | | | control - fibers | -0.128 | 0.317 | -0.403 | 1.000 | | |
| | | | control - tire wear | -0.137 | 0.317 | -0.434 | 1.000 | | |
| | | | control - PE | -0.069 | 0.317 | -0.218 | 1.000 | | |
| *P. verrucosa* | Gross photosynthesis | t1 | control - beach | -0.202 | 0.206 | -0.980 | 0.982 | 90 | lmer(scale(x)~treatment + (1\|col) |
| | | | control - fibers | 0.125 | 0.206 | 0.608 | 1.000 | | |
| | | | control - tire wear | -0.087 | 0.206 | **-0.424** | 1.000 | | |
| | | | control - PE | -0.598 | 0.206 | **-2.902** | **0.015** | | |
| | | t2 | control - beach | -0.262 | 0.297 | **-0.883** | 0.755 | 89 | lmer(scale(log(x+10)~treatment + (1\|col) |
| | | | control - fibers | -0.110 | 0.292 | **-0.378** | 0.755 | | |
| | | | control - tire wear | -0.386 | 0.292 | **-1.320** | 0.560 | | |
| | | | control - PE | -0.704 | 0.292 | **-2.409** | 0.064 | | |
| *S. pistillata* | Gross photosynthesis | t1 | control - beach | -0.289 | 0.279 | **-1.034** | 1.000 | 90 | lmer(scale(log(x+10)~treatment + (1\|col) |
| | | | control - fibers | -0.176 | 0.279 | **-0.629** | 1.000 | | |
| | | | control - tire wear | -0.273 | 0.279 | **-0.977** | 1.000 | | |
| | | | control - PE | -0.222 | 0.279 | **-0.793** | 1.000 | | |
| | | t2 | control - beach | -0.011 | 0.280 | **-0.039** | 1.000 | 90 | lmer(scale(log(x+10)~treatment + (1\|col) |
| | | | control - fibers | 0.062 | 0.280 | 0.222 | 1.000 | | |
| | | | control - tire wear | -0.150 | 0.280 | **-0.534** | 1.000 | | |
| | | | control - PE | -0.121 | 0.280 | **-0.430** | 1.000 | | |

Table S6: Results of statistical analyses assessing the effects of the microdebris treatments (beach, fibers, tire wear, and PE) compared to the control on the photosynthetic efficiency (i.e., effective quantum yield (Y(II)), maximum quantum yield (Fv/Fm), maximum relative electron transport rate ($rETR_{max}$), minimum saturating irradiance ($E_k$), and efficiency of light capture ($\alpha$)) of *Pocillopora verrucosa* and *Stylophora pistillata* after the first 4 weeks of the experiment (t1) and after 8 weeks of the experiment (t2). Results are derived from linear mixed effects models with treatment as fixed factor and colony (col) set as a random factor, on scale or log transformed data, followed by holm-adjustment for multiple testing. Numbers of observations (n), estimates, standard errors, z-values, and p-values are given with model specifications. Bold values indicate significance ($p < 0.05$).

| Species | Response variable | Time | Contrast | Estimate | Std. Error | z-value | p | n | Model specifications |
|---|---|---|---|---|---|---|---|---|---|
| *P. verrucosa* | Y(II) | t1 | control - beach | -0.470 | 0.236 | -1.993 | 0.185 | 90 | lmer(scale(log(x))~treatment + (1\|col)) |
| | | | control - fibers | -0.154 | 0.236 | -0.655 | 1.000 | | |
| | | | control - tire wear | -0.391 | 0.236 | -1.658 | 0.292 | | |
| | | | control - PE | 0.006 | 0.236 | 0.027 | 1.000 | | |
| | | t2 | control - beach | -0.354 | 0.134 | -2.643 | **0.025** | 90 | lmer(scale(x)~treatment + (1\|col)) |
| | | | control - fibers | -0.336 | 0.134 | -2.508 | **0.025** | | |
| | | | control - tire wear | -0.471 | 0.134 | -3.520 | **0.002** | | |
| | | | control - PE | 0.200 | 0.134 | 1.497 | 0.134 | | |
| *S. pistillata* | Y(II) | t1 | control - beach | -0.269 | 0.149 | -1.802 | 0.143 | 90 | lmer(scale(x)~treatment + (1\|col)) |
| | | | control - fibers | -0.426 | 0.149 | -2.855 | **0.013** | | |
| | | | control - tire wear | -0.637 | 0.149 | -4.266 | **0.000** | | |
| | | | control - PE | -0.063 | 0.149 | -0.423 | 0.672 | | |
| | | t2 | control - beach | -0.329 | 0.164 | -2.004 | 0.135 | 90 | lmer(scale(x)~treatment + (1\|col)) |
| | | | control - fibers | -0.072 | 0.164 | -0.438 | 0.923 | | |
| | | | control - tire wear | -0.601 | 0.164 | -3.662 | **0.001** | | |
| | | | control - PE | -0.121 | 0.164 | -0.736 | 0.923 | | |
| *P. verrucosa* | Fv/Fm | t1 | control - beach | -0.002 | 0.145 | -0.015 | 1.000 | 90 | lmer(scale(log(x))~treatment + (1\|col)) |
| | | | control - fibers | -0.010 | 0.145 | -0.068 | 1.000 | | |
| | | | control - tire wear | 0.113 | 0.145 | 0.777 | 1.000 | | |
| | | | control - PE | -0.040 | 0.145 | -0.277 | 1.000 | | |
| | | t2 | control - beach | -0.115 | 0.131 | -0.876 | 1.000 | 90 | lmer(scale(x)~treatment + (1\|col)) |
| | | | control - fibers | 0.011 | 0.131 | 0.085 | 1.000 | | |

| | | | | | | | | | | | |
|---|---|---|---|---|---|---|---|---|---|---|---|
| | | | control | - | tire wear | 0.083 | 0.131 | 0.632 | 1.000 | | |
| | | | control | - | PE | 0.002 | 0.131 | 0.012 | 1.000 | | |
| *S. pistillata* | Fv/Fm | t1 | control | - | beach | 0.141 | 0.139 | 1.015 | 0.620 | 90 | lmer(scale(x)~treatment + (1\|col)) |
| | | | control | - | fibers | 0.282 | 0.139 | 2.040 | 0.166 | | |
| | | | control | - | tire wear | 0.113 | 0.139 | 0.818 | 0.620 | | |
| | | | control | - | PE | 0.212 | 0.139 | 1.532 | 0.377 | | |
| | | t2 | control | - | beach | -0.090 | 0.091 | -0.991 | 0.965 | 90 | lmer(scale(x)~treatment + (1\|col)) |
| | | | control | - | fibers | -0.009 | 0.091 | -0.103 | 1.000 | | |
| | | | control | - | tire wear | -0.044 | 0.091 | -0.485 | 1.000 | | |
| | | | control | - | PE | -0.128 | 0.091 | -1.404 | 0.642 | | |
| *P. verrucosa* | $rETR_{max}$ | t1 | control | - | beach | -0.495 | 0.280 | -1.767 | 0.232 | 90 | lmer(scale(log(x))~treatment + (1\|col)) |
| | | | control | - | fibers | -0.394 | 0.280 | -1.404 | 0.245 | | |
| | | | control | - | tire wear | -0.621 | 0.280 | -2.216 | 0.107 | | |
| | | | control | - | PE | -0.433 | 0.280 | -1.545 | 0.245 | | |
| | | t2 | control | - | beach | -0.371 | 0.286 | -1.299 | 0.582 | 90 | lmer(scale(x)~treatment + (1\|col)) |
| | | | control | - | fibers | -0.904 | 0.286 | -3.164 | **0.006** | | |
| | | | control | - | tire wear | -0.224 | 0.286 | -0.782 | 0.868 | | |
| | | | control | - | PE | 0.023 | 0.286 | 0.079 | 0.937 | | |
| *S. pistillata* | $rETR_{max}$ | t1 | control | - | beach | -0.146 | 0.292 | -0.500 | 1.000 | 90 | lmer(scale(log(x))~treatment + (1\|col)) |
| | | | control | - | fibers | 0.465 | 0.292 | 1.592 | 0.446 | | |
| | | | control | - | tire wear | 0.042 | 0.292 | 0.144 | 1.000 | | |
| | | | control | - | PE | 0.164 | 0.292 | 0.562 | 1.000 | | |
| | | t2 | control | - | beach | 0.315 | 0.303 | 1.038 | 0.897 | 90 | lmer(scale(x)~treatment + (1\|col)) |
| | | | control | - | fibers | 0.392 | 0.303 | 1.293 | 0.785 | | |
| | | | control | - | tire wear | -0.171 | 0.303 | -0.565 | 1.000 | | |
| | | | control | - | PE | 0.057 | 0.303 | 0.189 | 1.000 | | |
| *P. verrucosa* | $E_k$ | t1 | control | - | beach | -0.382 | 0.247 | -1.545 | 0.355 | 90 | lmer(scale(log(x))~treatment + (1\|col)) |
| | | | control | - | fibers | -0.278 | 0.247 | -1.123 | 0.355 | | |
| | | | control | - | tire wear | -0.530 | 0.247 | -2.143 | 0.128 | | |
| | | | control | - | PE | -0.386 | 0.247 | -1.562 | 0.355 | | |

| | | | | | | | | | |
|---|---|---|---|---|---|---|---|---|---|
| | | t2 | control - beach | -0.219 | 0.264 | -0.827 | 1.000 | 90 | lmer(scale(x)~treatment + (1\|col)) |
| | | | control - fibers | -0.768 | 0.264 | -2.906 | **0.015** | | |
| | | | control - tire wear | -0.050 | 0.264 | -0.189 | 1.000 | | |
| | | | control - PE | 0.134 | 0.264 | 0.506 | 1.000 | | |
| *S. pistillata* | $E_k$ | t1 | control - beach | -0.172 | 0.291 | -0.590 | 1.000 | 90 | lmer(scale(log(x))~treatment + (1\|col)) |
| | | | control - fibers | 0.254 | 0.291 | 0.874 | 1.000 | | |
| | | | control - tire wear | -0.006 | 0.291 | -0.022 | 1.000 | | |
| | | | control - PE | -0.011 | 0.291 | -0.039 | 1.000 | | |
| | | t2 | control - beach | 0.305 | 0.277 | 1.102 | 1.000 | 90 | lmer(scale(log(x))~treatment + (1\|col)) |
| | | | control - fibers | 0.266 | 0.277 | 0.962 | 1.000 | | |
| | | | control - tire wear | -0.031 | 0.277 | -0.111 | 1.000 | | |
| | | | control - PE | 0.160 | 0.277 | 0.576 | 1.000 | | |
| *P. verrucosa* | α | t1 | control - beach | -0.361 | 0.223 | -1.620 | 0.421 | 90 | lmer(scale(x)~treatment + (1\|col)) |
| | | | control - fibers | -0.330 | 0.223 | -1.483 | 0.421 | | |
| | | | control - tire wear | -0.296 | 0.223 | -1.328 | 0.421 | | |
| | | | control - PE | -0.197 | 0.223 | -0.887 | 0.421 | | |
| | | t2 | control - beach | -0.421 | 0.186 | -2.259 | **0.048** | 90 | lmer(scale(x)~treatment + (1\|col)) |
| | | | control - fibers | -0.489 | 0.186 | -2.627 | **0.035** | | |
| | | | control - tire wear | -0.480 | 0.186 | -2.575 | **0.035** | | |
| | | | control - PE | -0.289 | 0.186 | -1.549 | 0.122 | | |
| *S. pistillata* | α | t1 | control - beach | -0.001 | 0.195 | -0.003 | 1.000 | 90 | lmer(scale(x)~treatment + (1\|col)) |
| | | | control - fibers | 0.439 | 0.195 | 2.254 | 0.097 | | |
| | | | control - tire wear | 0.093 | 0.195 | 0.478 | 1.000 | | |
| | | | control - PE | 0.321 | 0.195 | 1.647 | 0.299 | | |
| | | t2 | control - beach | 0.169 | 0.205 | 0.827 | 0.816 | 90 | lmer(scale(x)~treatment + (1\|col)) |
| | | | control - fibers | 0.274 | 0.205 | 1.340 | 0.668 | | |
| | | | control - tire wear | -0.283 | 0.205 | -1.382 | 0.668 | | |
| | | | control - PE | -0.155 | 0.205 | -0.758 | 0.816 | | |

Table S7: Results of PERMANOVA assessing the differences between the effects of the three microdebris mixtures (beach, fibers, and tire wear) and the single polymer (PE) on growth (i.e., surface area, volume, and calcification), photosynthesis rates (i.e., net photosynthesis, gross photosynthesis, and respiration) and the photosynthetic efficiency (i.e., effective quantum yield (Y(II)), maximum quantum yield (Fv/Fm), maximum relative electron transport rate ($rETR_{max}$), minimum saturating irradiance ($E_k$), and efficiency of light capture ($\alpha$)) from both measurement time points of the experiment (after 4 and 8 weeks). Results are derived from Permutational Multivariate Analysis of Variance with species set as strata. Degrees of freedom (Df), sum of squares (SS) $R^2$, F-value, and p-value are given.

| | Df | SS | $R^2$ | F | p |
|---|---|---|---|---|---|
| Treatment | 3 | 0.01766 | 0.00699 | 0.6617 | 0.638 |
| Residual | 282 | 2.50886 | 0.99301 | | |
| Total | 285 | 2.52652 | 1 | | |

Table S8: Results of statistical analyses assessing the effects of the three microdebris mixtures (beach, fibers, and tire wear) compared to the single polymer (PE) on growth (i.e., surface area, volume, and calcification), photosynthesis rates (i.e., net photosynthesis, gross photosynthesis, and respiration) and the photosynthetic efficiency (i.e., effective quantum yield (Y(II)), maximum quantum yield (Fv/Fm), maximum relative electron transport rate ($rETR_{max}$), minimum saturating irradiance ($E_k$), and efficiency of light capture ($\alpha$)) of *Pocillopora verrucosa* and *Stylophora pistillata* from both measurement time points of the experiment (after 4 and 8 weeks). Results are derived from linear mixed effects models with treatment as fixed factor and measurement timepoint (time) and colony (col) set as a random factor, on scale or log transformed data, followed by holm-adjustment for multiple testing. Numbers of observations (n), estimates, standard errors, z-values, and p-values are given with model specifications. Bold values indicate significance ($p < 0.05$).

<table>
<tr><th>Species</th><th>Response variable</th><th>Contrast</th><th>Estimate</th><th>Std. Error</th><th>z-value</th><th>p</th><th>n</th><th>Model specifications</th></tr>
<tr><td>*P. verrucosa*</td><td>Surface area</td><td>PE - beach</td><td>-0.002</td><td>0.139</td><td>0.016</td><td>1.000</td><td>179</td><td>lmer(scale(x)~treatment + (1|time) + (1|col))</td></tr>
<tr><td></td><td></td><td>PE - fibers</td><td>-0.274</td><td>0.138</td><td>1.982</td><td>0.142</td><td></td><td></td></tr>
<tr><td></td><td></td><td>PE - tire wear</td><td>0.028</td><td>0.138</td><td>-0.205</td><td>1.000</td><td></td><td></td></tr>
<tr><td>*S. pistillata*</td><td>Surface area</td><td>PE - beach</td><td>-0.021</td><td>0.125</td><td>0.170</td><td>1.000</td><td>180</td><td>lmer(scale(x)~treatment + (1|time) + (1|col))</td></tr>
<tr><td></td><td></td><td>PE - fibers</td><td>0.028</td><td>0.125</td><td>-0.220</td><td>1.000</td><td></td><td></td></tr>
<tr><td></td><td></td><td>PE - tire wear</td><td>0.001</td><td>0.125</td><td>-0.005</td><td>1.000</td><td></td><td></td></tr>
<tr><td>*P. verrucosa*</td><td>Volume</td><td>PE - beach</td><td>0.071</td><td>0.163</td><td>-0.437</td><td>1.000</td><td>179</td><td>lmer(scale(x)~treatment + (1|time) + (1|col))</td></tr>
<tr><td></td><td></td><td>PE - fibers</td><td>-0.275</td><td>0.162</td><td>1.700</td><td>0.268</td><td></td><td></td></tr>
<tr><td></td><td></td><td>PE - tire wear</td><td>-0.037</td><td>0.162</td><td>0.227</td><td>1.000</td><td></td><td></td></tr>
<tr><td>*S. pistillata*</td><td>Volume</td><td>PE - beach</td><td>0.165</td><td>0.139</td><td>-1.180</td><td>0.527</td><td>180</td><td>lmer(scale(x)~treatment + (1|time) + (1|col))</td></tr>
<tr><td></td><td></td><td>PE - fibers</td><td>0.189</td><td>0.139</td><td>-1.355</td><td>0.527</td><td></td><td></td></tr>
<tr><td></td><td></td><td>PE - tire wear</td><td>0.029</td><td>0.139</td><td>-0.208</td><td>0.835</td><td></td><td></td></tr>
<tr><td>*P. verrucosa*</td><td>Calcification</td><td>PE - beach</td><td>0.141</td><td>0.146</td><td>-0.965</td><td>1.000</td><td>180</td><td>lmer(scale(log(x+100)~treatment + (1|time) + (1|col))</td></tr>
<tr><td></td><td></td><td>PE - fibers</td><td>0.041</td><td>0.143</td><td>-0.284</td><td>1.000</td><td></td><td></td></tr>
<tr><td></td><td></td><td>PE - tire wear</td><td>0.003</td><td>0.148</td><td>-0.020</td><td>1.000</td><td></td><td></td></tr>
<tr><td>*S. pistillata*</td><td>Calcification</td><td>PE - beach</td><td>0.309</td><td>0.173</td><td>-1.788</td><td>0.221</td><td>178</td><td>lmer(scale(x)~treatment + (1|time) + (1|col))</td></tr>
<tr><td></td><td></td><td>PE - fibers</td><td>0.299</td><td>0.174</td><td>-1.716</td><td>0.221</td><td></td><td></td></tr>
<tr><td></td><td></td><td>PE - tire wear</td><td>0.043</td><td>0.173</td><td>-0.248</td><td>0.804</td><td></td><td></td></tr>
</table>

| | | | | | | | | |
|---|---|---|---|---|---|---|---|---|
| *P. verrucosa* | Net photosynthesis | PE - beach | 0.403 | 0.189 | -2.136 | **0.033** | 179 | lmer(scale(x)~treatment + (1\|time) + (1\|col)) |
| | | PE - fibers | 0.716 | 0.188 | -3.819 | **<0.001** | | |
| | | PE - tire wear | 0.610 | 0.188 | -3.252 | **0.002** | | |
| *S. pistillata* | Net photosynthesis | PE - beach | 0.023 | 0.190 | -0.120 | 1.000 | 180 | lmer(scale(x)~treatment + (1\|time) + (1\|col)) |
| | | PE - fibers | 0.166 | 0.190 | -0.872 | 1.000 | | |
| | | PE - tire wear | 0.000 | 0.190 | 0.001 | 1.000 | | |
| *P. verrucosa* | Respiration | PE - beach | 0.202 | 0.200 | -1.010 | 0.937 | 179 | lmer(scale(x)~treatment + (1\|time) + (1\|col)) |
| | | PE - fibers | 0.128 | 0.199 | -0.645 | 0.937 | | |
| | | PE - tire wear | -0.188 | 0.199 | 0.944 | 0.937 | | |
| *S. pistillata* | Respiration | PE - beach | 0.150 | 0.212 | -0.709 | 1.000 | 180 | lmer(scale(x)~treatment + (1\|time) + (1\|col)) |
| | | PE - fibers | -0.012 | 0.212 | 0.055 | 1.000 | | |
| | | PE - tire wear | 0.018 | 0.212 | -0.086 | 1.000 | | |
| *P. verrucosa* | Gross photosynthesis | PE - beach | 0.421 | 0.187 | -2.253 | **0.039** | 179 | lmer(scale(x)~treatment + (1\|time) + (1\|col)) |
| | | PE - fibers | 0.652 | 0.186 | -3.512 | **0.001** | | |
| | | PE - tire wear | 0.433 | 0.186 | -2.335 | **0.039** | | |
| *S. pistillata* | Gross photosynthesis | PE - beach | 0.093 | 0.209 | -0.447 | 1.000 | 180 | lmer(scale(x)~treatment + (1\|time) + (1\|col)) |
| | | PE - fibers | 0.128 | 0.209 | -0.616 | 1.000 | | |
| | | PE - tire wear | 0.009 | 0.209 | -0.043 | 1.000 | | |
| *P. verrucosa* | Effective quantum yield | PE - beach | -0.514 | 0.135 | 3.820 | **<0.001** | 180 | lmer(scale(x)~treatment + (1\|time) + (1\|col)) |
| | | PE - fibers | -0.354 | 0.135 | 2.626 | **0.009** | | |
| | | PE - tire wear | -0.547 | 0.135 | 4.064 | **<0.001** | | |
| *S. pistillata* | Effective quantum yield | PE - beach | -0.207 | 0.116 | 1.787 | 0.148 | 180 | lmer(scale(x)~treatment + (1\|time) + (1\|col)) |
| | | PE - fibers | -0.151 | 0.116 | 1.305 | 0.192 | | |
| | | PE - tire wear | -0.524 | 0.116 | 4.540 | **<0.001** | | |
| *P. verrucosa* | Maximum quantum yield | PE - beach | -0.045 | 0.100 | 0.451 | 1.000 | 180 | lmer(scale(x)~treatment + (1\|time) + (1\|col)) |
| | | PE - fibers | 0.018 | 0.100 | -0.186 | 1.000 | | |
| | | PE - tire wear | 0.113 | 0.100 | -1.131 | 0.774 | | |
| *S. pistillata* | Maximum quantum yield | PE - beach | -0.008 | 0.092 | 0.087 | 1.000 | 180 | lmer(scale(x)~treatment + (1\|time) + (1\|col)) |
| | | PE - fibers | 0.096 | 0.092 | -1.043 | 0.891 | | |
| | | PE - tire wear | 0.007 | 0.092 | -0.075 | 1.000 | | |

| | | | | | | | | |
|---|---|---|---|---|---|---|---|---|
| *P. verrucosa* | ETRmax | PE - beach | -0.196 | 0.205 | 0.958 | 0.415 | 180 | lmer(scale(x)~treatment + (1\|time) + (1\|col)) |
| | | PE - fibers | -0.472 | 0.205 | 2.302 | 0.064 | | |
| | | PE - tire wear | -0.258 | 0.205 | 1.260 | 0.415 | | |
| *S. pistillata* | ETRmax | PE - beach | -0.039 | 0.209 | 0.186 | 0.852 | 180 | lmer(scale(x)~treatment + (1\|time) + (1\|col)) |
| | | PE - fibers | 0.318 | 0.209 | -1.519 | 0.386 | | |
| | | PE - tire wear | -0.199 | 0.209 | 0.953 | 0.681 | | |
| *P. verrucosa* | Ek | PE - beach | -0.132 | 0.187 | 0.704 | 0.521 | 180 | lmer(scale(x)~treatment + (1\|time) + (1\|col)) |
| | | PE - fibers | -0.419 | 0.187 | 2.238 | 0.076 | | |
| | | PE - tire wear | -0.211 | 0.187 | 1.125 | 0.521 | | |
| *S. pistillata* | Ek | PE - beach | -0.050 | 0.199 | 0.252 | 1.000 | 180 | lmer(scale(x)~treatment + (1\|time) + (1\|col)) |
| | | PE - fibers | 0.163 | 0.199 | -0.817 | 1.000 | | |
| | | PE - tire wear | -0.164 | 0.199 | 0.822 | 1.000 | | |
| *P. verrucosa* | alpha | PE - beach | -0.148 | 0.143 | 1.031 | 0.736 | 180 | lmer(scale(x)~treatment + (1\|time) + (1\|col)) |
| | | PE - fibers | -0.167 | 0.143 | 1.162 | 0.736 | | |
| | | PE - tire wear | -0.144 | 0.143 | 1.008 | 0.736 | | |
| *S. pistillata* | alpha | PE - beach | 0.015 | 0.144 | -0.103 | 0.918 | 180 | lmer(scale(x)~treatment + (1\|time) + (1\|col)) |
| | | PE - fibers | 0.280 | 0.144 | -1.954 | 0.152 | | |
| | | PE - tire wear | -0.176 | 0.144 | 1.225 | 0.441 | | |

Table S9: Results of statistical analyses assessing the differences in polyp behavior (reaction, uptake, and egestion) of *Pocillopora verrucosa* and *Stylophora pistillata* between the particles of the microdebris treatments (beach, fibers, tire wear, and PE) and natural food (copepods). Results are derived from Fisher's exact tests (pairwise with holm correction for multiple testing) and stated as number of observations (n) and p-value. Bold values indicate significance ($p < 0.05$).

| Species | Observation | Contrast | | | n | p-value |
|---|---|---|---|---|---|---|
| *P. verrucosa* | Reactions | copepods | - | beach | 600 | **<0.001** |
| | | copepods | - | fibers | 600 | **<0.001** |
| | | copepods | - | tire wear | 600 | **<0.001** |
| | | copepods | - | PE | 600 | **<0.001** |
| | | beach | - | fibers | 600 | 0.019 |
| | | beach | - | tire wear | 600 | 0.822 |
| | | beach | - | PE | 600 | 0.342 |
| | | fibers | - | tire wear | 600 | **<0.001** |
| | | fibers | - | PE | 600 | **<0.001** |
| | | tire wear | - | PE | 600 | 0.822 |
| *S. pistillata* | | copepods | - | beach | 600 | **<0.001** |
| | | copepods | - | fibers | 600 | **<0.001** |
| | | copepods | - | tire wear | 600 | **<0.001** |
| | | copepods | - | PE | 600 | **<0.001** |
| | | beach | - | fibers | 600 | 1.000 |
| | | beach | - | tire wear | 600 | 1.000 |
| | | beach | - | PE | 600 | 0.696 |
| | | fibers | - | tire wear | 600 | 1.000 |
| | | fibers | - | PE | 600 | 0.181 |
| | | tire wear | - | PE | 600 | 0.254 |
| *P. verrucosa* | Uptake | copepods | | beach | 194 | **<0.001** |
| | | copepods | | fibers | 177 | **<0.001** |
| | | copepods | | tire wear | 201 | **<0.001** |
| | | copepods | | PE | 207 | **<0.001** |
| | | beach | | fibers | 35 | 1.000 |
| | | beach | | tire wear | 59 | 1.000 |
| | | beach | | PE | 65 | 1.000 |
| | | fibers | | tire wear | 42 | 1.000 |
| | | fibers | | PE | 48 | 1.000 |
| | | tire wear | | PE | 72 | 1.000 |
| *S. pistillata* | | copepods | | beach | 176 | **<0.001** |
| | | copepods | | fibers | 172 | **<0.001** |
| | | copepods | | tire wear | 173 | **<0.001** |
| | | copepods | | PE | 185 | **<0.001** |
| | | beach | | fibers | 24 | 1.000 |
| | | beach | | tire wear | 25 | 1.000 |
| | | beach | | PE | 37 | 1.000 |
| | | fibers | | tire wear | 21 | 1.000 |
| | | fibers | | PE | 33 | 1.000 |

| | | | | | |
|---|---|---|---|---|---|
| | | tire wear | PE | 34 | 1.000 |
| *P. verrucosa* | Egestion | copepods | beach | 154 | 1.000 |
| | | copepods | fibers | 154 | 1.000 |
| | | copepods | tire wear | 154 | 1.000 |
| | | copepods | PE | 157 | **0.002** |
| | | beach | fibers | 0 | 1.000 |
| | | beach | tire wear | 0 | 1.000 |
| | | beach | PE | 3 | 1.000 |
| | | fibers | tire wear | 0 | 1.000 |
| | | fibers | PE | 3 | 1.000 |
| | | tire wear | PE | 3 | 1.000 |
| *S. pistillata* | | copepods | beach | 128 | 1.000 |
| | | copepods | fibers | 129 | **0.078** |
| | | copepods | tire wear | 128 | 1.000 |
| | | copepods | PE | 128 | 1.000 |
| | | beach | fibers | 1 | 1.000 |
| | | beach | tire wear | 0 | 1.000 |
| | | beach | PE | 0 | 1.000 |
| | | fibers | tire wear | 1 | 1.000 |
| | | fibers | PE | 1 | 1.000 |
| | | tire wear | PE | 0 | 1.000 |

Table S10: Results of statistical analyses assessing the differences in interaction times of *Pocillopora verrucosa* and *Stylophora pistillata* between food and microdebris particles for positive reactions (reaction = yes), uptake events (uptake = yes), and rejection events (uptake = no). Results are derived from Wilcox tests and stated as the number of observations for food and microdebris particles (n), v-statistic, and p-value. Bold values indicate significance ($p < 0.05$).

| Species | Data | n food | n microdebris | v-statistic | p-value |
|---|---|---|---|---|---|
| *P. verrucosa* | Reaction = yes | 168 | 107 | 5212 | **<0.001** |
| *S. pistillata* | | 162 | 58 | 2274 | **<0.001** |
| *P. verrucosa* | Uptake = yes | 154 | 3 | 60 | **0.026** |
| *S. pistillata* | | 128 | 1 | 120 | 0.126 |
| *P. verrucosa* | Uptake = no | 14 | 104 | 600 | 0.288 |
| *S. pistillata* | | 34 | 57 | 442 | **<0.001** |

Table S11: Results of statistical analyses assessing the differences in polyp behavior (i.e., reactions, uptake, and egestion) between *Pocillopora verrucosa* and *Stylophora pistillata* to the different particles of the microdebris treatments (beach, fibers, tire wear, and PE) and natural food (i.e., copepods). Results are derived from Fisher's exact tests and stated as number of observations (n) and p-value. Bold values indicate significance ($p < 0.05$).

| Observation | Particle type | n | p |
|---|---|---|---|
| Reactions | Copepods | 600 | 0.682 |
| | Microdebris (all) | 2400 | **<0.001** |
| | Beach | 600 | 0.071 |
| | Fibers | 600 | 1 |
| | Tire wear | 600 | **0.001** |
| | PE | 600 | **0.044** |
| Uptake | Copepods | 330 | **0.002** |
| | Microdebris (all) | 165 | 1 |
| | Beach | NA | |
| | Fibers | 19 | 1 |
| | Tire wear | NA | |
| | PE | 62 | 0.288 |
